# ZBTB24 is a conserved multifaceted transcription factor at genes and centromeres that governs the DNA methylation state and expression of satellite repeats

**DOI:** 10.1101/2023.08.31.555516

**Authors:** Giacomo Grillo, Ekaterina Boyarchuk, Seed Mihic, Ivana Ivkovic, Mathilde Bertrand, Alice Jouneau, Thomas Dahlet, Michael Dumas, Michael Weber, Guillaume Velasco, Claire Francastel

**Affiliations:** UMR7216 Épigénétique et Destin Cellulaire, CNRS, Université de Paris Cité, Epigenetics and Cell Fate, F-75013, Paris, France; UMR7216, Genome engineering in epigenetics platform (GENIE); Bioinformatics and Biostatistics Core Facility, iCONICS, Institut du Cerveau (ICM), Sorbonne Université, INSERM, CNRS, Hôpital Pitié-Salpêtrière, Paris, France; Université Paris-Saclay, UVSQ, INRAE, BREED, 78350, Jouy-en-Josas, France; Ecole Nationale Vétérinaire d’Alfort, BREED, 94700, Maisons-Alfort, France; University of Strasbourg, Strasbourg, France. CNRS UMR7242, Biotechnology and Cell Signaling, Illkirch, France; Princess Margaret Cancer Centre, University Health Network, Toronto, Ontario, Canada

**Author notes:** Correspondence to: Guillaume Velasco, and Claire Francastel.

## Abstract

Since its discovery as an Immunodeficiency with Centromeric instability and Facial anomalies syndrome-causative gene, ZBTB24 has emerged as a key player in DNA methylation, immunity and development. By extensively analyzing ZBTB24 genomic functions in ICF-relevant mouse and human cellular models, we revealed here its multiple facets as a transcription factor, with key roles in immune response-related genes expression and also in early embryonic development. Using a constitutive *Zbtb24* ICF-like mutant and an auxin-inducible degron system in mouse embryonic stem cells, we showed that ZBTB24 is recruited to centromeric satellite DNA where it is required to establish the correct DNA methylation patterns through the recruitment of DNMT3B. Thus, our results further revealed an essential role for ZBTB24 at human and mouse centromeric satellite arrays, as a transcriptional repressor. Together, we unveiled unprecedented functions of ZBTB24 at human and mouse centromeres by directly controlling DNA methylation and transcription of the underlying tandem satellite repeats.

## INTRODUCTION

The study of diseases with abnormal DNA methylation (DNAme) landscapes emphasized the key role of this epigenetic mark in the maintenance of genomic stability (1–3). In particular, a growing body of evidence coming from studies of the etiology of these diseases, including but not limited to cancer, has suggested the existence of a functional link between CpG methylation and centromere integrity. Centromeres being chromosomal domains specialized in faithful segregation of the genetic material during cell divisions, which assemble on large-scale arrays of tandem repeated sequences, this functional link is thought to have a major contribution to genome stability (4, 5).

The inherited disorder called the ICF syndrome (Immunodeficiency with Centromeric instability and Facial anomalies syndrome; MIM #242860) is one of the most emblematic pathological contexts that has underlined the connection between DNAme and centromere integrity. This syndrome is a very rare autosomal recessive disorder, genetically heterogeneous, which is mostly characterized by primary immunodeficiency (6, 7). Although displaying varying degrees of severity, clinical manifestations also include mild facial anomalies, intellectual disability, congenital malformations and developmental delay (8). The ICF syndrome has been described as a disease affecting DNAme at constitutive heterochromatin domains assembled on juxtacentromeric satellite repeats: Satellites type II (Sat II) of chromosomes 1 & 16 and, to a lesser extent, Satellites type III (Sat III) of chromosome 9. DNA hypomethylation of pericentromeric satellite DNA is an invariant hallmark of the ICF patients’ cells, which leads to juxtacentromeric heterochromatin decondensation, chromosomal breaks and rearrangements in the vicinity of centromeres, a molecular feature used to establish diagnosis (9). In contrast, additional hypomethylation of centromeric alpha satellite DNA is a heterogeneous trait among ICF patients, thus reflecting the genetic heterogeneity of the syndrome (10).

The ICF syndrome is caused by homozygote or heterozygote composite mutations in the DNA methyltransferase 3B gene (*DNMT3B*, ICF1 MIM#242860) (11, 12), zinc finger and BTB domain containing 24 gene (*ZBTB24*, ICF2 MIM#614069) (13), cell division cycle-associated 7 gene (*CDCA7*, ICF3 MIM#616910) (14), or helicase lymphoid specific gene (*HELLS*, ICF4 MIM#616911) (14). Strikingly, only DNMT3B has a DNA methyltransferase (DNMT) activity whereas the other factors are transcription factors (TFs) (ZBTB24 and CDCA7) or chromatin remodelers (HELLS/LSH) with poorly known functions, questioning their role in DNAme pathways. Our comparative methylome analysis of ICF patients revealed a striking similarity between genome-wide DNAme profiles of ICF2-4 patients, which clearly distinguished them from ICF1 patients (15, 16). Remarkably, ICF2-4 patient subtypes also exhibit hypomethylation of centromeric satellite repeats in addition to that of the juxtacentromeric satellites common to all ICF subtypes (10, 15). Thus, the ICF syndrome represents a unique pathological context whose study has allowed to identify unanticipated molecular actors and pathways linking DNAme to centromere integrity.

In contrast to DNMT3B, the functional connections between ZBTB24, CDCA7 and HELLS with the DNA methylation machinery are not straightforward but begin to be clarified. Indeed, ZBTB24 was shown to be a transcriptional activator of the *CDCA7* gene (17), whereas CDCA7 is required for HELLS to associate with chromatin and stimulates its remodeling activity (18). As supported by recent studies, a CDCA7/HELLS complex contributes to DNAme maintenance by facilitating the recruitment of the DNMT1/UHRF1 complex (19, 20). In light of these data, and combined to our previous study showing that ZBTB24 function is required to maintain DNAme at centromeric repeats in mouse embryonic fibroblasts (14), a molecular model has been proposed along which the ZBTB24-CDCA7 axis may regulate HELLS functions in DNAme maintenance of centromeric DNA repeats in murine cells (21). However, the direct contribution of ZBTB24 to the DNAme process, especially during the DNAme establishment step and at these repeated sequences, remains untested.

ZBTB24 belongs to the broad-complex, tramtrack and bric-à-brac - zinc finger (BTB-ZF) family of transcription factors known for their essential role in the development of the immune system (22). ZBTB24 is constitutively expressed, with the highest expression in B cells (13), suggesting an important role in the function of this cell lineage. ZBTB24 harbors a BTB domain thought to mediate protein oligomerization and/or dimerization, an AT-hook motif through which it can bind to AT-rich DNA sequences and eight Zinc-finger motifs in its C-terminal part, supporting the fact that ZBTB24 is a transcription factor (22). Indeed, as shown by studies using a tagged version of the protein in the HCT116 colon cancer cell line or in mouse embryonic stem cells (mESCs), exogenous ZBTB24 acts as a transcriptional regulator, mostly as an activator, which localizes at gene regulatory regions and recognizes a specific DNA motif conserved between mouse and human (23, 24). Interestingly, in murine NIH3T3 cells, exogenous ZBTB24 protein localizes to chromocenters, i.e. subnuclear clusters of (peri)centromeres from several chromosomes, suggesting that ZBTB24 would also play an important role at (peri)centromeric heterochromatin (25, 26). As shown in zebrafish, ZBTB24 function would be essential for the methylated status of pericentromeric satellite DNA repeats and their transcriptional repression (27).

Considering that its loss of function is the cause of the ICF2 syndrome, we explored here the potential direct contribution of ZBTB24 functions to the immune response pathways, embryonic development, DNAme and centromere integrity. Taking advantage of mouse and human cellular systems and models relevant for the physiopathological features of the ICF syndrome, combined to multi-omics approaches, we addressed the multiple facets of the endogenous ZBTB24 protein. Notably, we document the genomic occupancy of endogenous ZBTB24 at regulatory sequences of genes and the transcriptomic consequences of its loss of function in human lymphocytes cells and in a dynamic cell-state transition during the differentiation process of mouse embryonic stem cells. In addition, we unveiled the striking and conserved occupancy of ZBTB24 at centromeric DNA repeats in both mouse and human, despite the poor conservation of centromeric sequences among species. We found that ZBTB24 has distinct roles at unique genes and tandem repeats with completely distinct genomic organization, mainly as a trans-activator of gene expression or a repressor of centromeric transcription. Importantly, our data indicate that ZBTB24 controls centromeric repeats transcription through the regulation of their DNAme status and the recruitment of the DNA methyltransferase DNMT3B. As a whole, this study highlights the dual role played by ZBTB24 depending on genomic location and provides new insights into its contribution to the maintenance of centromeric integrity, which alterations invariantly lead to chromosomal instability diseases.

## RESULTS

### ZBTB24 is a regulator of innate response-related genes in human lymphocytes

To explore the role of ZBTB24 as a transcription factor and the consequences of its pathological loss of function (LOF) on the transcriptome, we performed chromatin immunoprecipitation of endogenous ZBTB24 followed by DNA sequencing (ChIP-seq) in human lymphoblastoid cell lines (LCLs) from three healthy donors (HDs) and RNA-sequencing (RNA-seq) in LCLs derived from HDs and two ICF2 patients (**Fig. S1A**). We chose this cellular context because i) it is particularly relevant to the pathophysiology of the ICF syndrome and ii) the ZBTB24 protein is expressed at high levels in LCLs compared to other somatic cell lines (13).

ChIP-seq data analysis revealed 183 genomic sites occupied by ZBTB24 in LCLs derived from HDs (**Table S1**, merge peaks, q-value <0.05), which predominantly localized at transcription start sites (TSS) or within proximal promoters (≤1Kb) and, to a lesser extent, at distal intergenic regions (**Fig.1A and 1B**). The functional annotation of ZBTB24 sites, according to the chromHMM genome segmentation ENCODE data in LCLs (GM12878), showed that it mainly occupies active promoters (n=106, “Active_Promoter” category) and enhancers (n=32, “Strong_Enhancer” category) (**Fig.1A**). ZBTB24 peaks showed a highly significant enrichment of the 5’-CAGGACCT-3’ DNA motif (E_value=2.8e-34) (**Fig. 1A**, lower panel), identical to the one identified for a FLAG-tagged ZBTB24 protein in the colon cancer cell line HCT116 (23).

**Figure 1.**
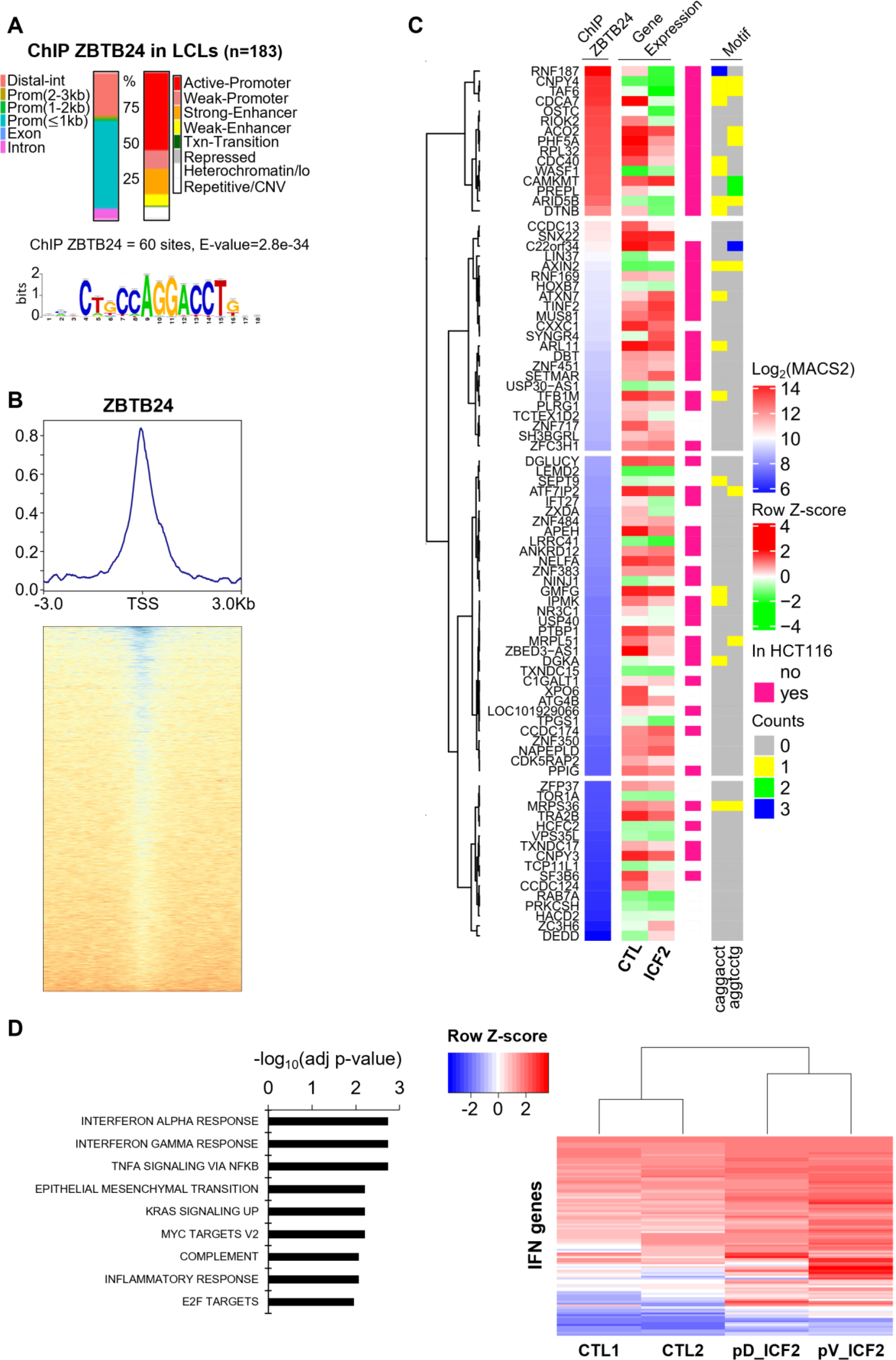
ZBTB24 acts as transcriptional regulator modulating innate immune response-related genes in patients-derived LCLs. (A) ChIP-seq analyses showing the distribution of genomic sites occupied by ZBTB24 (183 peaks) in lymphoblastoid cell lines (LCLs) across genomic (left) and functional annotations (right). The functional annotation was based on the chromatin states established in GM12878 LCLs (ChromHMM track from ENCODE/Broad Institute). Predicted binding motif of ZBTB24 was identified using Regulatory Sequence Analysis Tools (RSAT, http://rsat.sb-roscoff.fr/). Distal_int, Distal intergenic; Prom, promoter. (B) Profile of the average of ZBTB24 ChIP-seq signal, *i.e* the enrichment of ZBTB24 relative to chromatin input (substraction mode, input from IP of bigWig files) in LCLs on genomic loci defined as 3kb upstream and downstream of annotated Transcriptional Start Site (TSS, NCBI RefSeq) of genes (upper panel). The lower panel shows the heatmap of read density around the same genomic loci. High density levels are shown in blue and low-density levels in red. (C) Unsupervised clustering of genes TSS linked to ZBTB24 sites. The heatmap (ChIP ZBTB24) presents the enrichment of ZBTB24 at each TSS based on the peak score (mean log_2_(MACS2 score)). The heatmap also shows the gene expression data, presented as a mean of the row Z-score of each gene in LCLs derived from the blood of healthy (CTL) and ICF2 subjects (ICF2). ZBTB24 genomic sites common to LCLs and HCT116 cell line (GSE111689, Thompson et al 2018) are notified in pink. The Motif columns refer to the occurrence of the predicted ZBTB24 binding motif occurrence (Counts). The color code attributed to each category is depicted next right to the figure. (D) Gene Set Enrichment Analysis (GSEA) of deregulated genes in LCLs derived from the blood of ICF2 patients (pD and pV) (left panel). The expression status of genes belonging to the “Interferon_response” HALLMARK category is presented as the row Z-score and illustrated by a heatmap (right panel).

Based on the peak score (MACS2 score) at TSS, we identified the main target genes of ZBTB24 which are *CDCA7, RIOK2, CNPY4, TAF6, ARID5B, CAMKMT, PREPL, RNF187, RPL32, CDC40, WASF1, OSTC, ACO2, PHF5A, and DTNB* (**Fig.1C**). Notably, for most of them, their expression levels were lower in LCLs from ICF2 patients compared to HDs, indicating that ZBTB24 is a transcriptional activator for this set of genes (**Fig.1C**). These results, combined with those previously reported (23), strongly suggest that ZBTB24 preferentially targets the TSS of genes in a sequence specific manner to positively regulate their expression.

However, the whole transcriptome analysis of ICF2 LCLs identified 356 up- and 50 down-regulated genes compared to normal LCLs (log2FC >0.5, FDR <0.01), suggesting a role of ZBTB24 as a negative gene regulator with a significant enrichment in a set of genes involved in the interferon response (adjusted p-value= 1.82e-03, **Table S2 and Fig.1D**) and, more broadly, in immune response pathways (**Fig. S1B**), although at these particular genes it may be an indirect consequence.

As a whole, these results show that, in human LCLs, ZBTB24 deficiency is associated with overexpression of genes involved in immune response pathways, more specifically of innate immune response-related genes, which is consistent with the autoimmune signs in the ICF patients (28, 29).

### ZBTB24 is essential for mouse embryonic development

The molecular and phenotypic features of ICF2 patients clearly pinpoint a critical role for ZBTB24 during development since nearly all patients exhibit developmental delay and growth retardation (8). To better define this role, we generated a mouse model for the ICF2 syndrome that reproduced the BTB domain mutations reported in Lebanese siblings (30) (**Fig. S1A and S2A-S2D**). Of note, the ZBTB24 protein sequence is highly conserved between mouse and human ranging from 76.9 to 96.9 % of homology for the structural domains (26).

Crosses of mice heterozygous for ZBTB24 mutation (*Zbtb24^wt/mt^*) did not produce newborns homozygous for the mutation (*Zbtb24^mt/mt^*), indicating the embryonic lethality of *Zbtb24^mt/mt^* embryos (**Table S3**). We isolated embryos at different developmental stages [6.5, 9.5- and 11.5-days post coitum (dpc)] and noticed that *Zbtb24^mt/mt^* embryos had a growth delay at all stages and presented morphological abnormalities (**Fig.2A**). These observations highlighted an essential role for ZBTB24 during early mouse development, consistent with the growth delay that characterizes ICF patients.

**Figure 2.**
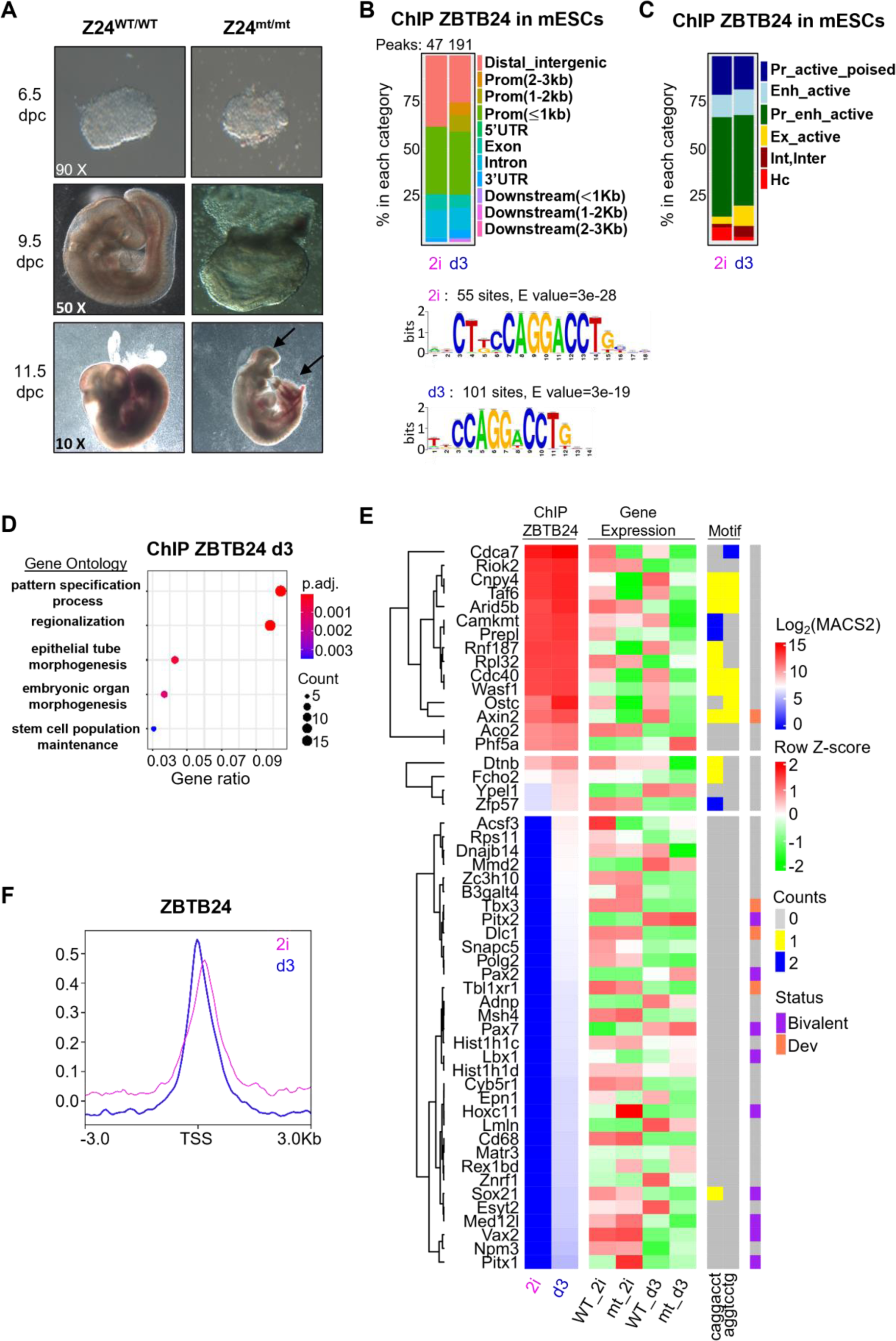
ZBTB24 targets developmental genes in mESCs and is essential for embryonic development. (A) Pictures of *Zbtb24^wt/wt^*, *Zbtb24^wt/mt^* and *Zbtb24^mt/mt^* embryos at post-implantation stages 6.5, 9.5- and 11.5-days post coitum (dpc). The arrows point to the embryonic defects reported in *Zbtb24^mt/mt^* embryos at 11.5 dpc. (B) Stacked bar chart of the distribution of the genomic sites occupied by ZBTB24 across genomic features. Peaks overlapping between two independent mESC clones derived from distinct embryos cultured in 2i conditions (2i) or at day3 of differentiation (d3) were merged and annotated to the mm10 genome. The predicted ZBTB24 binding motif identified by Regulatory Sequence Analysis Tools (RSAT, http://rsat.sb-roscoff.fr/) based on ChIP-seq data performed in 2i condition (2i) or at day3 of differentiation (d3) is reported below the plots. (C) Stacked bar chart of the distribution of ZBTB24 sites across functional regions of the mouse genome (mm10). ZBTB24 peaks overlapping between two independent ES clones cultured in 2i conditions (2i) or at day3 of differentiation (d3) were merged and annotated based on ChIP-seq assays of 8 histone modifications as part of the ENCODE 3 Consortium (laboratory of Bing Ren). Pr: promoter; Enh: enhancer; Int,Inter: intron, intergenic; Ex: exon; Hc: heterochromatin. (D) Gene Ontology of genes linked to ZBTB24 sites in mESCs at day3 of differentiation (ChIP ZBTB24 d3). The dot plot represents the enrichment in biological processes. GeneRatio on the x axis refers to the number of genes linked to ZBTB24 sites divided by the number of genes annotated in the corresponding process. The size and colors of the dots indicate gene counts and the adjusted p-value for the GO pathway listed on the y axis, respectively. Only the top 5 GO terms with the highest degree of enrichment are represented (GO:0007389 pattern specification process, GO:0003002 regionalization, GO:0060562 epithelial tube morphogenesis, GO:0048562 embryonic organ morphogenesis, GO:0019827 stem cell population maintenance). (E) Unsupervised clustering of genes linked to ZBTB24 merge peaks at TSS. The heatmap (ChIP ZBTB24) presents the enrichment of ZBTB24 at TSS of genes based on the peak score (mean log_2_(MACS2 score)), in two independent ES clones cultured in 2i conditions (2i) or at day3 of differentiation (d3). For each locus, the gene status (“Bivalent” or “Dev.”: developmental genes) and the predicted ZBTB24 binding motif occurrence are indicated, and a color code is attributed to each category next right to the figure. The gene expression data are presented as the mean of the row Z-score in the two independent ES clones WT and mutant (mt) for ZBTB24 cultured in 2i condition (WT_2i, mt_2i) or at day3 of differentiation (WT_d3, mt_d3). (F) Average of ChIP_seq signal profile of ZBTB24, *i.e* the enrichment of ZBTB24 relative to chromatin input (substraction mode, input from IP) in mESCs cultured in 2i conditions (2i) or at day3 of differentiation (d3) on genomic loci defined as 3kb upstream and downstream of annotated Transcriptional Start Site (TSS, NCBI RefSeq) of genes.

To better explore this role, we first derived and characterized mouse embryonic stem cells (mESCs) from wild-type (*Zbtb24^wt/wt^*) and mutant (*Zbtb24^mt/mt^*) blastocysts, isolated at 3.5 dpc (31), in 2i medium (two inhibitors of the MAPK/ERK and GSK3β pathways) to maintain their naïve ground state (32) (**Fig.S2E-S2F**). We took advantage of the dynamic cell-state transition during the differentiation process of mESCs to model the first molecular steps that occur in the early embryo. We performed ChIP-seq of endogenous ZBTB24 in *Zbtb24^wt/wt^*mESCs and RNA-seq in *Zbtb24^wt/wt^* and *Zbtb24^mt/mt^* mESCs maintained in 2i conditions (2i) or after 3 days of differentiation (d3). While the levels of ZBTB24 protein in *Zbtb24^wt/wt^*mESCs were nearly identical in 2i and at d3 (**Fig.S2G**), we identified a four times greater number of ZBTB24 genomic binding sites in differentiated mESCs (191 sites, q-value <0.05) compared to mESCs in 2i (47 sites, q-value <0.05) (**Fig. 2B** and **Table S4-S5**). Nearly all the ZBTB24 sites identified in 2i were maintained in differentiated mESCs (44 out of 47 peaks). At d3, ZBTB24 binding sites localized mainly at cis-regulatory elements, active promoters and enhancers (**Fig.2B and 2C**).

The top 5 GO terms of genes linked to ZBTB24 peaks in differentiated mESCs, but not in 2i, matched remarkably with the developmental defects reported in the *Zbtb24^mt/mt^*embryos and are all linked to parent biological processes “Anatomical structure morphogenesis” and “Multicellular organismal process” (**Fig. 2D** and **Table S6**). Nonetheless, most of the genes belonging to these GO terms and categorized as developmental or bivalent (**Fig. 2E**, based on ENCODE3 data), showed relatively weak ZBTB24 peaks consistent with the absence of the identified DNA binding motif, suggesting they are not primary targets of ZBTB24 at this early stage of differentiation. This could be a reason why we did not detect major transcriptional deregulation of this set of genes in *Zbtb24^mt/mt^* mESCs cellular model at d3 of differentiation (**Fig. 2E**).

Together, these results showed that ZBTB24 is essential for mouse embryonic development possibly through its ability to target and regulate some key developmental genes.

### ZBTB24 is a conserved transcriptional activator of a set of unique genes

We next evaluated the conservation of the role of ZBTB24 as a transcriptional regulator in mice and humans (**Fig.1 & 2**). Compared to human cells, we identified the same functional and chromatin state features for ZBTB24 genomic sites in mESCs, an identical enriched DNA motif 5’-CAGGACCT-3’within ZBTB24 peaks and a conserved genomic distribution, *i.e*. at promoters and at distal intergenic regions (**Fig.2B**), at TSS (21/47 peaks in 2i and 55/191 peaks at d3, **Fig.2F**) and enhancers, mostly at d3 (3 peaks in 2i compared to 35 peaks at d3) (**Fig.2C**). Noteworthy, most of the genes showing the highest ZBTB24 enrichment at TSS were identical to those identified in human LCLs (*Cdca7, Riok2, Cnpy4, Taf6, Arid5b, Camkmt, Prepl, Rnf187, Rpl32, Cdc40, Wasf1, Ostc, Axin2, Aco2, Phf5a and Dtnb*) (**Fig.S3**), suggesting similar mechanisms of regulation of gene expression in humans and mice. These target genes exhibited reduced expression levels in *Zbtb24^mt/mt^* mESCs, strongly supporting a conserved role of ZBTB24 as a transcriptional activator at this conserved set of unique genes (**Fig.2E and Fig.S3**).

As a whole, these results revealed a discrete and conserved role of ZBTB24 as a transcriptional activator at unique genes, both in humans and mice. These results also validated the use of our ICF2 mouse model to further decipher the conserved genomic functions of ZBTB24.

### ZBTB24 plays a minor role in CpGme establishment at genes and interspersed repeats

Considering the central role played by DNAme during mammalian development (33) and the altered DNAme patterns that are typical of ICF2 patient cells (15), we further explored the direct role of ZBTB24 in shaping DNAme patterns during early embryonic development.

We have previously reported that ZBTB24 is required for the maintenance of DNAme at mouse centromeric DNA repeats (14), along with a widespread DNA hypomethylation in ICF2 patients (15). To address the role of ZBTB24 in DNAme establishment, which occurs in the early phases of mouse embryonic development, we profiled the DNA methylome of *Zbtb24^wt/wt^*and *Zbtb24^mt/mt^* mESCs, in naïve and differentiated state (d3), using Reduced Representation Bisulfite Sequencing (RRBS). Indeed, the dynamic mESCs transition from 2i to d3 is recognized as a powerful model that mimics *in vitro* the embryonic window where DNAme profiles are established *de novo* (34, 35). As expected, the DNAme profiles obtained by RRBS allowed to clearly distinguish mESCs in 2i medium from those at d3 of differentiation, with a gain of both weakly and strongly methylated sequences in differentiated mESCs (**Fig.3A and 3B**).

**Figure 3.**
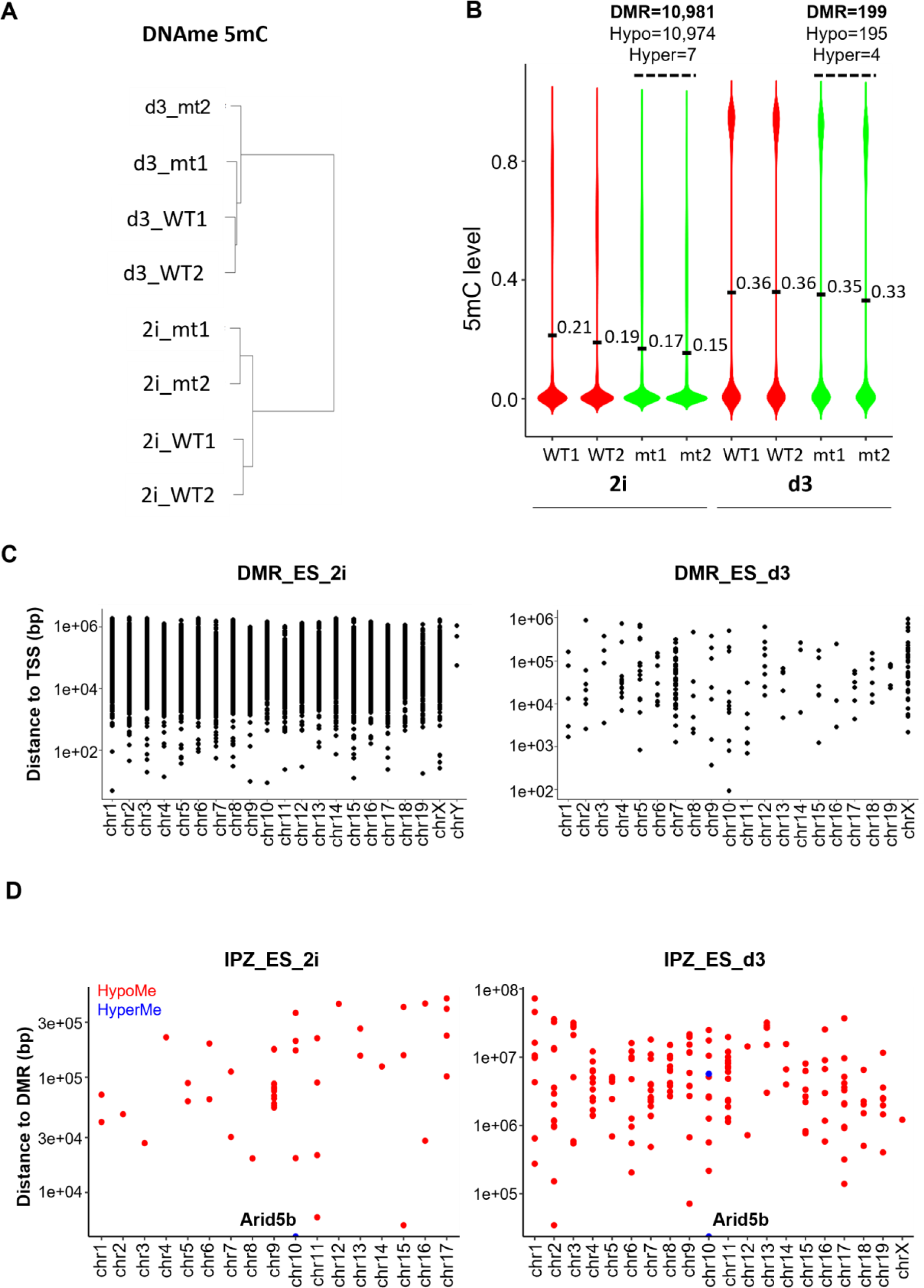
ZBTB24 moderately contributes to the establishment of DNAme landscape. (A) Dendrogram showing the clustering of the two mESCs clones wild-type (WT) and mutant (mt) for ZBTB24, cultured in 2i medium (2i) or at day3 of differentiation (d3), based on their DNAme profiles performed by RRBS. (B) Violin plot of DNAme level in the two ES clones wild-type (WT) and mutant (mt) for ZBTB24, cultured in 2i medium or at day3 of differentiation. The number and the status (Hypo: Hypomethylation; Hyper: Hypermethylation) of Differentially Methylated Regions (DMR) identified in ZBTB24 mutant compared to WT mESCs is shown next above to the plot. (C) Dot plots showing the distance in base pairs (bp) between DMR and TSS of genes in mESCs cultured in 2i medium or at day 3 of differentiation. (D) Dot plots presenting the distance in base pairs (bp) between ZBTB24 peak summits in WT mESCs and DMR (hypomethylated in red, hypermethylated in blue) in WT mESCs cultured in 2i medium and at day3 of differentiation (IPZ_ES_2i and IPZ_ES_d3). Dots which correspond to genes whose change in their level of expression is correlated to DMR position are indicated by gene symbol.

In naïve and differentiated mESCs, *Zbtb24^mt/mt^* cells showed distinct DNAme profiles compared to *Zbtb24^wt/wt^* mESCs, which indicates that ZBTB24 is required to ensure correct shaping of DNAme profiles at early development stages (**Fig.3A**). Surprisingly, we observed reduced levels of DNAme in *Zbtb24^mt/mt^* compared to *Zbtb24^wt/wt^*mESCs in 2i conditions (10,974 hypomethylated methylated regions or hypoDMR) (**Fig.3B and Table S7**), a state at which the levels of DNAme are known to be low and minimal compared to mESCs cultured in serum conditions (35, 36). Upon commitment to differentiation, the number of hypoDMR in *Zbtb24^mt/mt^* cells was greatly reduced compared to *Zbtb24^wt/wt^* cells (195 *vs* 10,974 hypoDMR), suggesting that the wave of DNAme establishment that occurs during this process compensated most of the methylation defects identified in the naive state despite the ZBTB24 LOF (**Fig.3B and Table S8**). HypoDMRs were mostly annotated away (>1kb) from TSS (**Fig.3C**), i.e. to intergenic regions, within LINE (Long Interspersed Nuclear Elements) and ERV (Endogenous Retroviruses) retrotransposon elements (**Fig.S4A-S4B and Table S7-S8**). However, hypomethylation of these retrotransposon elements was not accompanied by a change in their expression (**Fig. S4C**).

To assess whether the methylation status of the genomic sites occupied by ZBTB24 was directly impacted by ZBTB24 LOF, we computed the distance between ZBTB24 sites (peak summits) and the position of the DMRs identified in *Zbtb24^mt/mt^*mESCs. We applied the same analyses to human cells using our previous methylome profiling in ICF2 patients (15). In both cases, we found that hypoDMRs in ZBTB24 mutant cells were located far away (>10Kb) from ZBTB24 binding sites both in mouse and human cells (**Fig.3D and supplementary Fig.S4D**). Notably, we identified a restricted number of hypermethylated (hyperMe) sites, mostly detectable in human LCLs, that were closer (<1Kb) to ZBTB24 genomic targets compared to the hypoMe ones. Interestingly, these hyperMe sites resided within the promoter region of *RNF187*, *CDC40*, *ZNF717,* and *ARID5B* which are ZBTB24 target genes and whose expression levels were downregulated in ICF2 patient cells (**Fig.S3 and Fig.1C**). Of note, the hyperMe of CpG within the promoter of *ARID5B* and its downregulation in ZBTB24 mutant cells was a conserved feature shared by human and mouse cells mutant for ZBTB24. (**Fig.3D**).

Altogether, as inferred from ZBTB24 LOF, ZBTB24 does not seem to be directly involved in the DNAme establishment step for a large proportion of the genome. However, our methylome highlights potential role of ZBTB24 in the protection of some of its main genomic targets against DNAme gain during development (37).

### ZBTB24 targets centromeric DNA repeats and maintains them in a silent state

The identification of *ZBTB24* mutations as a genetic cause of the ICF syndrome naturally raised the question of its role in centromere DNAme and integrity. Given the presence of an AT-hook within ZBTB24 structural domains, we hypothesized that ZBTB24 was able to directly target centromeric satellite sequences. To test this hypothesis, we overcame the problem posed by the high sequence homology between satellite repeats units and their tandemly repeated nature, by using the Repenrich2 method (38) to quantify the enrichment of ZBTB24 ChIP-seq reads in centromeric sequences within the regions occupied by ZBTB24 (see Materials and Methods section). Of note, the genome organization of mouse and human centromeres are clearly distinct and not conserved. Mouse centromere DNA repeats are relatively homogeneous between all chromosomes and composed of AT-rich tandem repeats of 120 bp called minor satellite (MinSat), annotated as SYNREP_MM and CENSAT_MC in the Repeatmasker database (**Fig.S5A and S5B**). Mouse pericentromeric DNA is composed of tandem repeats of 234 bp major satellite sequences annotated as GSAT_MM in the Repeatmasker database. In contrast to mouse centromeres, human centromeres are composed of Alpha satellite DNA, monomers of 171 bp arranged in a head-to-tail manner and organized in repeated arrays called higher order repeats (HORs) that are chromosome-specific (39).

Based on our ZBTB24 ChIP-seq data obtained from *Zbtb24^wt/wt^*mESCs, we identified reads mapping to centromeric satellite DNA with a modest enrichment in differentiated mESCs (d3) compared to 2i conditions (**Fig.4A**). These analyses were further confirmed by ChIP-qPCR using primers specific for MinSat repeats (**Fig.4B**). Using the same approach in human cells, we were able to identify ChIP-seq reads that matched ALR-Alpha satellite sequences, and also the pericentromeric satellite repeats HSAT4, HSATII, SAR, SATR1, and SATR2 (**Fig.4C**). Altogether, our results strongly indicate that ZBTB24 occupies centromeric satellite DNA in mouse and human cells.

**Figure 4.**
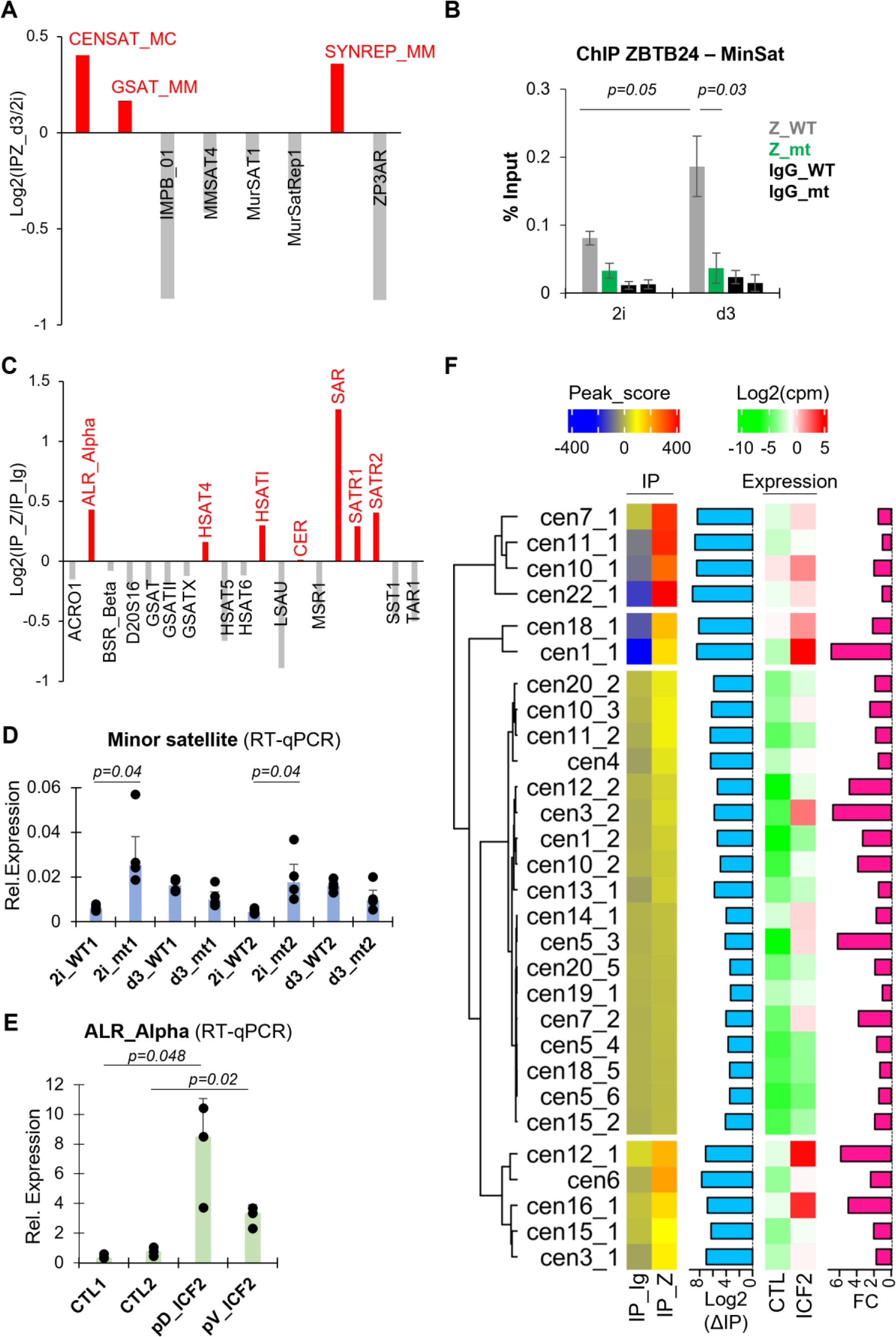
ZBTB24 occupies centromeric satellite DNA and regulates cenRNA level. (A) Bar chart showing the enrichment of normalized read counts in mouse satellite repeats based on ZBTB24 occupancy in mESCs at day3 of differentiation compared to 2i condition. The satellite repeats which show an enrichment of ZBTB24 occupancy in differentiated mESCs are shown in red and labeled. (B) ChIP assays performed on chromatin prepared from WT(Z_WT) and ZBTB24 mutant (Z_mt) mESCs cultured in 2i medium or at day 3 of differentiation, using ZBTB24 antibodies or IgG, and followed by quantitative PCR with primers specific of Minor satellite sequences. Error bars represent standard error (n=3 independent experiments), p : p-value; two-tailed t-test (C) Bar chart showing the enrichment of normalized read counts in human satellite repeats based on ChIP-seq of ZBTB24 and of IgG in LCLs. The human satellite repeats which show an enrichment of ZBTB24 occupancy are shown in red and labelled. (D) Expression levels of Minor satellite RNA in WT and *Zbtb24* mutant (mt) mESCs cells clones (1 and 2) cultured in 2i medium or at day 3 of differentiation were assessed by RT-qPCR, presented as a relative expression to U6 snRNA levels. Error bars represent standard error (n=4 independent experiments), p : p-value; two-tailed t-test . (E) Expression levels of ALR_Alpha satellite RNA in LCLs derived from healthy (CTL1 and CTL2; WT for ZBTB24) and ICF2 subjects (pD, PV; mutant for ZBTB24) were assessed by RT-qPCR, presented as a relative expression to U6 snRNA levels. Error bars represent standard error (n=3 independent experiments), p : p-value; two-tailed t-test. (F) Unsupervised clustering of centromeric HORs occupied by ZBTB24. The heatmap (IP) presents the peak score computed from the mean difference between normalized reads count in ChIP_seq of ZBTB24 (IP_Z) and of IgG (IP_Ig) at the considered HOR in LCLs derived from the blood of three healthy subjects. The bar chart on the right shows the log2 difference of the peak score between IP_Z and IP_Ig. The heatmap (Expression) shows the expression data of each HOR computed from RNA-seq data of two independent LCL clones WT (CTL) and mutant (ICF2) for ZBTB24 derived from the blood of subjects. The bar chart on the right presents the log2 fold change (FC) of HORs expression based on RNA-seq data, comparing the ICF2 to the WT subjects.

Because of its role as a transcription factor, we tested whether ZBTB24 LOF impacted the levels of centromeric transcripts (cenRNA) originating from these centromeric regions. In mESCs grown in 2i, the levels of MinSat transcripts are low but significantly higher in the context of ZBTB24 LOF (**Fig.4D**). In human cells, ICF2 patient cells showed higher levels of Alpha satellite RNA (**Fig.4E**). These data strongly suggest that ZBTB24 is required to control the levels of centromeric transcripts.

We then performed deeper analyses of our sequencing reads mapping to human centromeres to investigate whether ZBTB24 occupies specific HORs and regulates the levels of transcripts generated from these HORs. We found that ZBTB24 occupancy was enriched at specific centromeric HORs with a marked preference for cen1_1, cen6, cen7_1, cen8, cen9, cen10_1, cen11_1, cen16_1, cen18_1, cen22_1 and cenX (**Fig. S6**). Interestingly, targeted HORs were mainly the active HORs, HOR_1, defined as such based on their enrichment in CENP-A and binding sites for CENP-B (CENP-Box) (40), two essential proteins for centromere function. It follows that the centromeric sites occupied by ZBTB24 are those enriched in CENP-B Box DNA binding motifs (**Fig. S6**). Furthermore, the levels of transcripts originating from HORs occupied by ZBTB24 were higher in ICF2 patient cells (**Fig.4F**).

Altogether, these results indicate that ZBTB24 preferentially occupy active HORs and mediate their silencing, likely through its ability to ensure proper DNA methylation at these satellite sequences.

### ZBTB24 is required for DNMT3B-mediated DNAme establishment at centromeres

Hypomethylation of centromeric satellite DNA is a molecular feature of ICF2 patient cells, and our previous data provided evidence for a crucial role of ZBTB24 in controlling the DNAme status of these repeated DNA sequences (**Fig.S7A and S7B**) (13–15). One of the most powerful techniques commonly used to assess DNAme status at Satellite DNA repeats is the Southern analysis using methyl-sensitive restriction enzymes. Southern blot data were represented and summarized as density and quantification plots of the radiolabeled blot signals (**Fig. 5A, 5B, 6C, 6F, S7B, S9A, S10B**) explained in **Fig.S8**.

**Figure 5.**
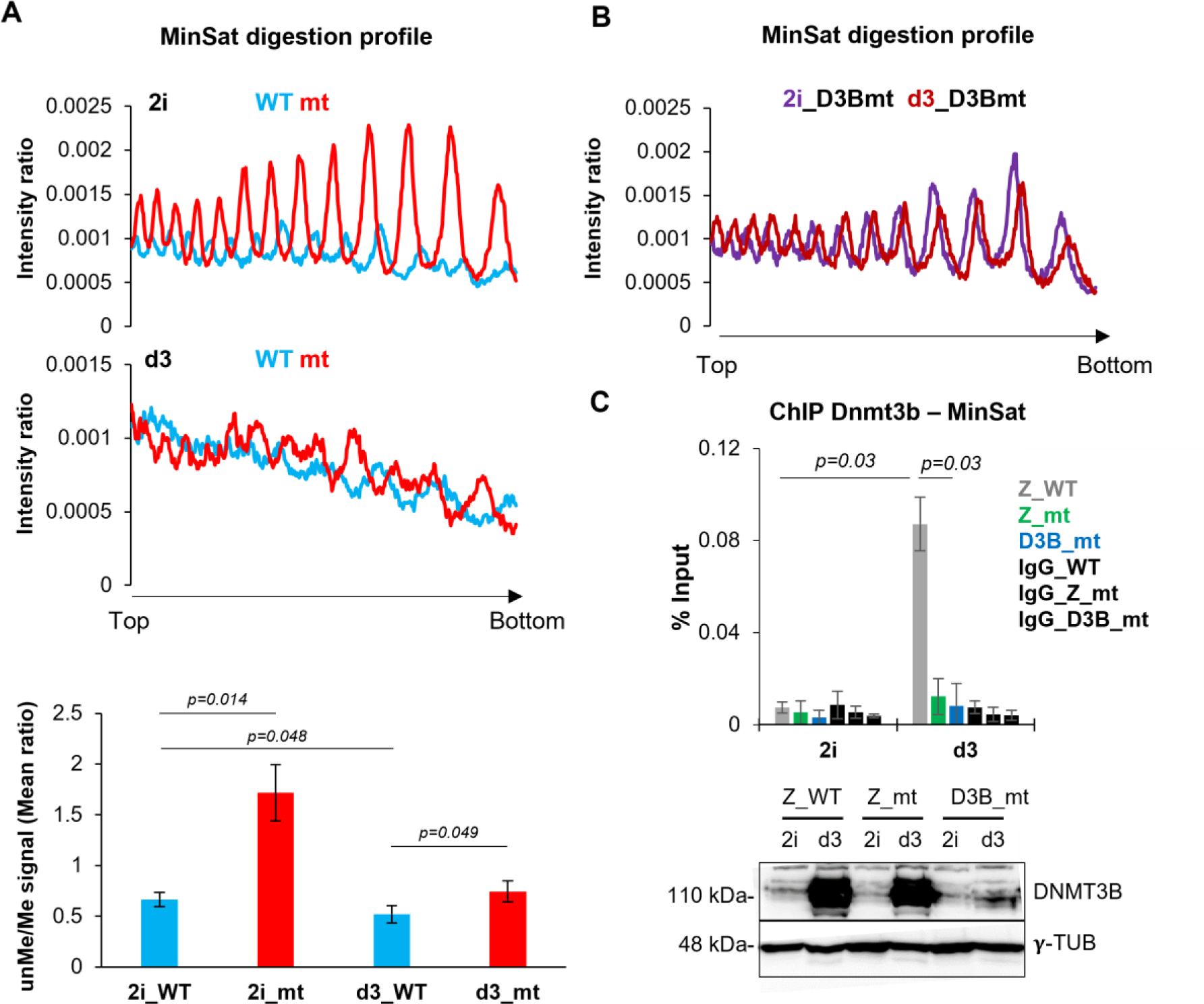
ZBTB24 is required for DNAme of MinSat during ES cells differentiation through DNMT3B recruitment. (A) Density plots of Southern blot band profiles obtained from the digestion of genomic DNA from *Zbtb24^wt/wt^* and *Zbtb24^mt/mt^*mESCs cultured in 2i medium (2i, top) and after 3 days of differentiation (d3, bottom) by the methylation-sensitive enzyme HpaII. The plot represents the profile of the radiolabeled signals intensity detected on each lane of the Southern blot after hybridization by radiolabeled minor satellite probes and normalized by the sum of the total signal per lane (from the top toward the bottom of the lane). The quantification of DNA methylation changes at minor satellite sequence is shown below the density plots. Error bars represent standard error (n=3 independent experiments on two different clones), p : p-value; two-tailed t-test (B) Density plots of Southern blot band profiles obtained from the digestion of genomic DNA from *Dnmt3b^mt/mt^*mESCs (D3Bmt) cultured in 2i medium or after 3 days of differentiation by the methylation-sensitive enzyme HpaII. The plot represents the profile of the radiolabeled signals intensity detected on each lane of the southern blot after hybridization by radiolabeled minor satellite probes and normalized by the sum of the total signal per lane (from the top toward the bottom of the lane). (C) (top) ChIP assays performed on chromatin prepared from *Zbtb24^wt/wt^*, *Zbtb24^mt/mt^* and DNMT3B mutant (*Dnmt3b^mt/mt^*) mESCs cultured in 2i medium or at day 3 of differentiation, using anti-ZBTB24, anti-DNMT3B or IgG antibodies, followed by quantitative PCR with primers specific of minor satellite sequence. Error bars represent standard error (n=3 independent experiments on two different clones), p : p-value; two-tailed t-test. (bottom) Western blot analysis showing the expression level of DNMT3B and γ-Tubulin proteins in *Zbtb24^wt/wt^*, *Zbtb24^mt/mt^* and *Dnmt3b^mt/mt^* mESCs cultured in 2i medium and at day3 of differentiation. Molecular weight is indicated on the left of the panel.

**Figure 6.**
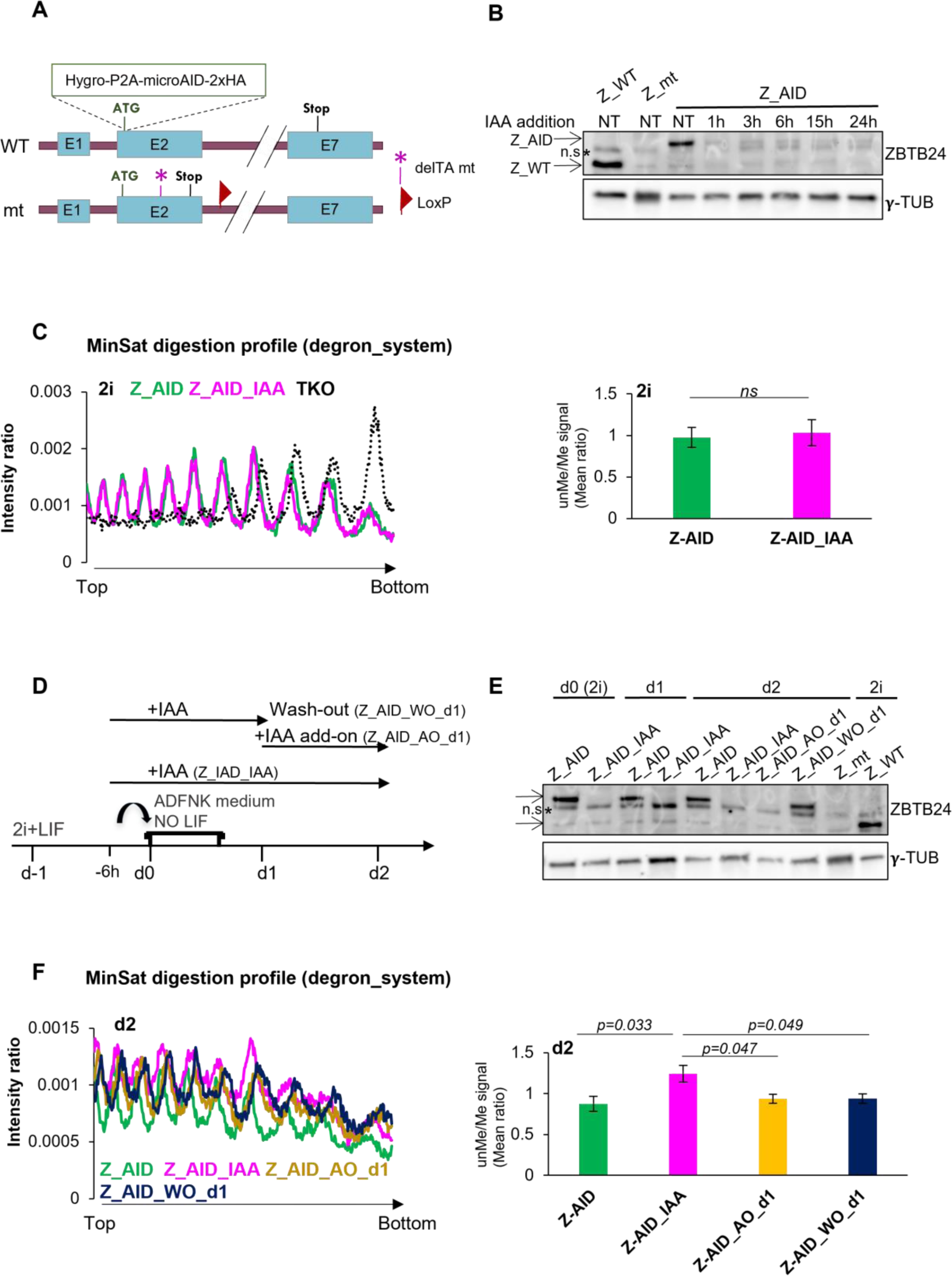
ZBTB24 is required for the early step of DNAme establishment of MinSat. A) Schematic representation of CRISPR/Cas9-based tagging of WT allele of ZBTB24 protein at N-terminus in heterozygous ZBTB24 ^wt/mt^ mESCs, expressing TIR1 from Rcc1 locus, to generate AID_ZBTB24 mESCs. Mutated *Zbtb24* allele with the position of the loxP site and the mutation is also shown. (B) Western blot analysis of ZBTB24 and γ-Tubulin protein levels in 2i condition in WT (Z_WT), mutant (Z_mt), and AID_ZBTB24 (Z_AID) cell lines. AID_ZBTB24 cells were treated with Auxin (IAA) for the indicated time. NT – non-treated with IAA, n.s. non-specific band. (C) (left panel) Density plot of Southern blot band profiles obtained from the digestion of genomic DNA, by the methylation-sensitive enzyme HpaII, of mESCs AID_ZBTB24 lines treated (Z_AID_IAA) or not (Z_AID) with auxin (IAA) cultured in 2i. The DNA from mESCs triple knocked-out (TKO) for the DNMTs enzymes was used as a control of the hypomethylated context. The quantification of DNA methylation changes at minor satellite sequence is shown on the right next to the density plots. Error bars represent standard error (n=3 independent experiments on two different clones), p : p-value; two-tailed t-test. (D) Experimental scheme for panels E and F. AID_ZBTB24 cells were either differentiated in ADFNK medium without LIF for 2 days (Z_AID) or treated with auxin (IAA) for the indicated time. For Z_ IAA and Z_IAA_wash-out, auxin was added 6h prior the differentiation induction. During the two days of mESCs differentiation, several conditions were used: auxin was either added 6h before the beginning of the differentiation protocol (Z_AID_IAA), at d1 (Z_AID_AO_d1) or remove at d1 (Z_AID_WO_d1). (E) Western blot analysis of ZBTB24 and γ-Tubulin protein levels in AID_ZBTB24 cells differentiated according to the schematic in D. Non-differentiated WT (Z_WT) and mutant (Z_mt) mESCs were used as controls, n.s. non-specific band. (F) Density plot of Southern blot band profiles obtained from the digestion of genomic DNA, by the methylation-sensitive enzyme HpaII, of mESCs AID_ZBTB24 lines treated (Z_AID_IAA) or not (Z_AID) with auxin (IAA), cultured in 2i and at day 1 or day 2 of differentiation (d1, d2). Auxin was either added 6h before the beginning of the differentiation protocol (Z_AID_IAA), at d1 (Z_AID_AO_d1) or removed at d1 (Z_AID_WO_d1). The plot represents the profile of the radiolabeled signals intensity detected on each lane of the Southern blot after hybridization by radiolabeled minor satellite probes and normalized by the sum of the total signal per lane (from the top toward the bottom of the lane). The quantification of DNA methylation changes at minor satellite sequence is shown on the right next to the density plots. Error bars represent standard error (n=3 independent experiments on two different clones), p : p-value; two-tailed t-test. Only significant differences are indicated.

To assess whether such a ZBTB24 function is conserved in mice, we analyzed the DNAme status of centromeric satellite sequences in *Zbtb24^wt/wt^*and *Zbtb24^mt/mt^* embryos, by Southern blot using DNAme-sensitive restriction enzymes. We found that MinSat repeated sequences were hypomethylated in *Zbtb24^mt/mt^* embryos, as evidenced by the signal intensity profiles reflecting the presence of low molecular weight DNA fragments that are produced by enzymatic digestion (**Fig.S9A and S9B**). These data showed that ZBTB24 is required for the methylated status of centromeric repeats in both mouse and human. Although we provided evidence that ZBTB24 is required for the maintenance of DNAme at centromeric MinSat repeats in murine somatic cells (14), the question of its involvement in the establishment of DNAme at these satellite repeats during early development remained unaddressed. Hence, we followed the DNAme status of MinSat during the wave of methylation that occurs as mESCs are induced to differentiation, using DNAme-sensitive restriction enzymes and Southern blotting. In agreement with our RRBS analysis (**Fig.3A and 3B**), we observed that MinSat were already strongly hypomethylated in *Zbtb24^mt/mt^* compared to *Zbtb24^wt/wt^* mESCs in 2i conditions (**Fig.5A**). This hypomethylated state was maintained in *Zbtb24^mt/mt^* cells at d3 of differentiation, although less pronounced than in 2i conditions, suggesting an incomplete or delayed establishment of DNAme patterns at MinSat repeats (**Fig.5A**).

DNA methyltransferase DNMT3B is not only required but is also the sole DNMT that catalyzes the *de novo* DNAme of MinSat sequences (41). Mouse ES cells mutant for DNMT3B (D3Bmt) failed to properly establish DNAme of MinSat at d3 of differentiation (**Fig.5B and S9C**). Hence, we tested whether ZBTB24 was involved in DNAme establishment at MinSat repeats through recruitment of DNMT3B to MinSat repeats. Whereas the expression levels of the DNMT3B protein was comparable between *Zbtb24^wt/wt^* and *Zbtb24^mt/mt^*mESCs (**Fig.5C**), ChIP-qPCR experiments clearly showed that there was a significant loss of DNMT3B occupancy at MinSat repeats in ZBTB24 LOF conditions (**Fig.5C**). Since the level of DNAme at MinSat in *Zbtb24^mt/mt^* mESCs was markedly lower than in *Zbtb24^wt/wt^* mESCs in 2i condition (**Fig.5A**), it is possible that the lower level of DNAme at MinSat observed in differentiated mutant cells (d3) could be due to the pre-existing hypomethylated status of these repeats in *Zbtb24^mt/mt^*compared to *Zbtb24^wt/wt^* mESCs in the naïve state.

In order to formally address the role of ZBTB24 in DNAme establishment at MinSat in an unbiased manner, we generated and characterized an auxin inducible degron (AID) knock-in WT mESCs lines (Z_AID) (**Fig.6A**) to specifically trigger ZBTB24 degradation at specific time points during mESCs differentiation. To validate the degradation of ZBTB24 upon auxin treatment in these Z-AID mESCs lines, we followed the levels of the ZBTB24 protein by western blot analysis at different time points, as well as the expression level of *Cdca7,* one of the major ZBTB24 target genes. We tested the impact on DNAme at MinSat at different times points of Auxin treatment or during the recovery dynamics after auxin wash-out. In 2i conditions, and after only 1h of auxin treatment, the levels of ZBTB24 protein dropped drastically whereas *Cdca7* expression significantly diminished 6h after auxin addition (**Fig. 6B and S10A**). After 24h and 48h of auxin treatment in 2i conditions, levels of DNAme were unchanged compared to untreated cells (**Fig.6C and S10B**), meaning that this auxin degron system will allow to formally compare the DNAme establishment kinetics at MinSat in cells expressing or not ZBTB24 since the DNAme level in naïve state is the same unlike the constitutive mutant of ZBTB24.

We then tested ZBTB24 requirement for DNAme establishment at MinSat repeats, using kinetics experiments to either degrade ZBTB24 protein in the presence of auxin (IAA Indole-3-acetic acid and Add On/AO conditions) or re-express ZBTB24 (Wash Out/WO) at specific time points during mESCs differentiation (**Fig.6D and 6E**). Differentiating mESCs (d2) in the presence of auxin failed to correctly establish DNAme patterns at MinSat, which strongly supports the crucial role of ZBTB24 during this process (**Fig.6F**). Moreover, the induced degradation of ZBTB24 at day 1 of differentiation (AO_d1) was sufficient to trigger failure of DNAme at MinSat. Rescuing ZBTB24 expression at day 1 of differentiation (WO_d1) was inefficient in fully restoring DNAme levels at MinSat (**Fig.6F**). These results indicated that ZBTB24 is required all along the process of DNAme establishment at MinSat sequences.

In sum, our results demonstrate an essential role of ZBTB24 in DNA methylation establishment at MinSat through guiding DNMT3B to these centromeric repeated sequences.

## DISCUSSION

ZBTB24 has long been considered as a ZBTB protein with an “obscure physiological function” (22), until the identification of mutations in *ZBTB24* as a causative factor of the ICF2 syndrome (13) brought to light its importance for development, the immune system, centromere biology and DNAme. Yet, being a transactivator of the *CDCA7* gene, ZBTB24 and CDCA7 were thought to be redundant in regulating gene expression, as it may be the case for genes linked to the immune response. Here, using mouse and human cellular systems relevant to ZBTB24 LOF that affect patients, combined to multi-omics approaches, we provide a list of conserved and direct targets of ZBTB24, which strikingly includes regulatory regions of a set of unique genes as well as tandemly repeated sequences that underlie centromeric chromosomal regions. At murine and human centromeres occupied by ZBTB24, we revealed that ZBTB24 is required for their transcriptional repression, and, to our knowledge, the first of its kind to be identified. We demonstrate that ZBTB24 is also required for the establishment of DNAme at centromeric repeats, through the recruitment of the *de novo* DNA methyltransferase DNMT3B. In combination with our demonstration of its role in DNAme maintenance at these loci in somatic cells (14), our study lend support to a model where ZBTB24 has pivotal and direct roles in the maintenance of centromere integrity, which alterations lead to chromosomal instability and disease.

### ZBTB24 LOF affects innate immune response-related genes

We reported here the first transcriptomic profiling of ICF2 patients-derived lymphocytes, which highlighted the deregulation of genes involved in signaling pathways linked to the innate immune response, as evidenced by an activation of genes involved in the interferon response. This supports the implication of ZBTB24 in cellular defense mechanisms, a function of the protein that is also supported by other studies reporting an activation of interferon response-related genes, upon knockdown of *ZBTB24* in the human B-cell line Raji and the colorectal cell line HCT116, or in a zebrafish ICF2 model (23, 27, 42). Since these interferon response related genes are not directly bound by ZBTB24, it suggests that their up-regulation is most likely an indirect consequence of ZBTB24 LOF. Of note, autoimmune symptoms have been reported for some ICF2 patients (28, 29), and it is worth mentioning that the activation of the interferon response pathways could account for this clinical trait. Interestingly, ZBTB24 LOF in zebrafish leads to an aberrant derepression of pericentromeric transcripts that triggers an interferon response in a zebrafish ICF2 model (27). Combined to the abnormally high levels of centromeric transcripts that we documented in the LCLs derived from ICF2 patients, it is tempting to speculate that these transcripts could trigger RNA surveillance mechanisms and, in turn, an antiviral response-like mediated by an activation of interferon response-related genes identified in ICF2 patient cells.

### ZBTB24 is essential for a healthy development in mouse and human

In mice, we showed that ZBTB24 is essential for early embryonic development. This result is consistent with a previous study using a mouse model carrying a larger genetic deletion of the BTB domain, AT-Hook and the first zinc fingers of ZBTB24 (17). Using mutations that recapitulate those found in ICF2 patients to create a mouse model for the disease and their derived mESCs, we identified a set of developmental genes occupied by ZBTB24 in differentiated mESCs whose functions are related to the growth delay and morphological abnormalities in *Zbtb24^mt/mt^* embryos.

In addition, in light of the pivotal role of DNAme in embryonic development (33) and given the deleterious impact of ZBTB24 LOF on the DNA methylome of patients (15) and murine cells (discussed below), it is likely that these alterations of DNAme landscapes could also account for the developmental defects observed in *Zbtb24^mt/mt^*embryos (this study) and human patients (28, 29).

The requirement of ZBTB24 for normal embryonic development seems to be more widely conserved during evolution since ZBTB24 mutations in zebrafish lead to early developmental abnormalities and lethality in adulthood (27), or to a developmental disease in humans. In addition, a strong decrease in *ZBTB24* expression is associated with recurrent spontaneous abortion in humans (43). Overall, ZBTB24 LOF leads to major developmental alterations, strongly supporting a conserved role for this protein during the early stages of development.

### Distinct roles of ZBTB24 in shaping DNA methylation patterns

We found that the genome of *Zbtb24^mt/mt^* naïve mESCs was remarkably hypomethylated compared to their wild-type counterpart in which DNAme levels are already minimal (35, 36). Certain classes of DNA repeats remain methylated in pluripotent mESCs as they escape the wave of demethyation after fertilization (44). Remarkably, among the sequences that loose DNA methylation upon ZBTB24 LOF, we found a number of these DNA repeats like LINE and ERVs. Hence, although we cannot exclude that ZBTB24 is required for DNA methylation establishment in primordial germ cells, ZBTB24 appears as being required for DNAme maintenance at about 10,000 genomic loci. Since ZBTB24 binds regulatory sequences at motifs that do not contain CG dinucleotide, it is unlikely that it has the ability to “read” and then protect the DNAme mark from demethylation waves, unlike ZFP57, another zinc-finger protein implicated in DNAme maintenance at imprinted loci (45, 46). In addition, although some other members of the BTB-ZF TF family are DNAme readers (47–49), the subnuclear localization of exogenous ZBTB24 at chromocenters in murine cells devoid of DNAme (25) suggest that more sophisticated mechanisms for DNA methylation maintenance are involved.

By mimicking “in-dish” the establishment of DNAme patterns in differentiating mESCs, we were able to show a rather modest contribution of ZBTB24 in this genome-wide process, reinforcing the idea that ZBTB24 plays mainly a role in DNAme maintenance. This observation is in line with previous studies showing progressive loss of DNAme at (peri)centromeric DNA repeats in HEK293 cells upon *ZBTB24* knock-out (50) or between larval and adult stages in *Zbtb24* mutant zebrafish (27) or else, DNA hypomethylation in HCT116 cells upon *ZBTB24* knockdown (23). It is not known whether DNAme defects in ICF2 patients worsen over time and whether this phenomenon correlates with an exacerbation of clinical signs as reported and suggested in a case report (29).

Importantly, we have dissected the role of ZBTB24 in DNAme of centromeric satellite sequences. On its own, the hypomethylated status of MinSat repeats in *Zbtb24^mt/mt^* embryos is not sufficient to conclude whether DNAme establishment, maintenance or both were altered upon loss of ZBTB24 function. Although we previously reported the requirement of ZBTB24 in DNAme maintenance at centromeric repeats in mouse embryonic fibroblasts (14), its contribution to DNAme establishment was not yet been assessed. Taking advantage of two genetic mESCs knock-in ICF2 models (*Zbtb24^mt/m^* ZBTB24-AID), we provided strong evidence that ZBTB24 is required for the proper establishment of DNAme at centromeric MinSat repeats. Moreover, we demonstrated that ZBTB24 has a direct role through its ability to bind to MinSat repeats and recruit DNMT3B. In line with this, a functional link between ZBTB24 and DNMT3B proteins at gene bodies has been reported in HEK 293 cells, also suggesting a role of ZBTB24 in DNAme establishment at these sequences (23). However, and because this interaction has not been confirmed in mESCs (21), we cannot rule out the possibility that the recruitment of DNMT3B to MinSat repeats is indirect. In a context of a crosstalk between DNA and histone methylation, as it was documented for H3K36me2/me3 and H3K9me2/me3 (51–53), DNMT3B recruitment may depend on a chromatin context shaped by the presence of ZBTB24, allowing the recruitment of the DNAme machinery. Yet, ZBTB24 is among the very rare TFs that have been identified as necessary for the localization of the DNAme machinery at specific places on the genome, here centromeric satellite sequences (54–56).

### ZBTB24 is a guardian of centromere integrity

Hence, we present here a novel facet of ZBTB24 function at centromeres. Our data provide strong evidence for a conserved and direct role for ZBTB24 at murine and human centromeric satellite sequences. Since these A/T rich sequences lack the DNA recognition motif of ZBTB24 found at regulatory sequences, read through its 6th and 7th zinc finger motifs (24), it is likely that the binding of ZBTB24 involves its AT-hook domain, which is known to interact with the minor groove of A/T rich DNA. However, as the AT-hook domain alone has a low affinity for DNA, the affinity of ZBTB24 for centromeric satellite sequences would be potentiated by the joint action of both AT-hook and ZnF motifs (57). In line with this, the presence of both motifs has been shown to be important for the localization of ZBTB24 to (peri)centromeric heterochromatin (26). Considering the growing body of evidence that points to a role of DNAme in centromeric integrity (4) and the fundamental role of centromeres in ensuring faithful transmission of the genetic information (58), ZBTB24 emerges as a guardian of centromeric integrity and hence, of genomic stability. In this study, we showed that the loss of DNAme at centromeric repeats upon ZBTB24 LOF was associated with increased levels of centromeric transcripts, more prominently in human cells, which have been shown to lead to chromosomal instability (59–62). Actually, the loss of DNAme at centromeric sequences could lead to their abnormal transcription, which poses a threat to genomic integrity through the formation of DNA:RNA hybrids and R-loops structures, known to cause replication stress or double strand breaks leading to genomic instability (50, 60, 61, 63–66). Interestingly, our in-depth analysis of the ChIP-seq data of ZBTB24 in human LCLs highlighted its peculiar enrichment at specific centromeric HORs, mainly the active ones (HOR-1) according to their enrichment in CENP-A and binding sites for CENP-B (CENP-Box) (40). Among the HORs mostly enriched in ZBTB24 occupancy and with deregulated expression in ICF2 cells, the presence of HOR_1 from chromosomes 1 and 16, which account for chromosomal anomalies in ICF patients, was striking. Whether the deregulation of specific HORs transcription in ICF cells is due to different degrees of DNAme loss or detectability due to a wide range of centromeres sizes in humans (67), and whether hypomethylation at pericentromeric regions of chromosomes 1 and 16 facilitates hypomethylation at nearby centromeric HORs are very exciting questions. These points should be addressed in a near future and be facilitated by the achievement of the complete sequence of the human genome, including centromeric satellite arrays, by the Telomere to Telomere (T2T) consortium (68).

Initially, the identification of three new ICFs factors lacking DNA methyltransferase activity, namely ZBTB24, CDCA7 and HELLS, raised the obvious question of their functional link with the DNAme machinery and centromere biology. Likely because of the known role of HELLS as “a guardian of heterochromatin at repeat elements” in mouse (69), and CDCA7 being downstream of ZBTB24 (17), most of the studies designed so far focused on the role of the CDCA7/HELLS complex (18, 19, 21, 50).

In sum, our work now provides a novel and previously unconsidered dimension of the role of ZBTB24 in maintaining genomic integrity.

## MATERIALS AND METHODS

### ICF2 mouse model

Mice mutant for *Zbtb24* were generated at the Institut Clinique de la Souris (Illkirch, France, http://www.ics-mci.fr) by homologous recombination using standard gene targeting techniques. The targeting construct was designed to replace exon 2 by mutated sequences (deletion of two nucleotides TA), thus mimicking the human mutations of ICF2 patients of a Lebanese family described in (30) carrying the deletion c.396_397delTA (p.His132Glnfs*19). Homozygous mice for *Zbtb24* mutation (*Zbtb24^mt/mt^*) were generated by intercrossing heterozygous mice (*Zbtb24^wt/mt^*) in a C57BL/6J background. Mice were mated overnight and the presence of a vaginal plug the following morning was designated as E0.5. Generation and use of these mice has been approved by the Comité d’éthique en Expérimentation Animale Buffon (Paris, France) and received the agreement number 5314. The genotyping was performed using genomic DNA extracted either from cultured cells, yolk sac or tail of embryos with the REDExtract-N-Amp Tissue PCR kit (Merck) following the manufacturer’s instructions.

### Cell lines and culture conditions

Mouse ES cells (mESCs) were derived from WT, *Zbtb24* and *Dnmt3b* (*Dnmt3b^m3/m3^*; Velasco, Hube et al., PNAS, 2010) mutant blastocysts as described in (31) at 3.5 dpc. ES cells deficient for all three active DNA methyltransferases, Dnmt1/Dnmt3a/Dnmt3b triple knockout (TKO), were kindly provided by Masaki Okano (70). All mESCs used in this study were maintained in the absence of feeder cells, on plates coated with 1x Laminin (Merck) as described in (71), in serum-free 2i medium containing Dulbecco’s modified Eagle’s medium (DMEM) F12 Glutamax (Thermo Fisher Scientific) supplemented with 50% Neurobasal medium (Thermo Fisher Scientific), 1x N2 and B27 Supplements (both Thermo Fisher Scientific), 1% penicillin/streptomycin (Merck), 1x non-essential amino acids (PAA), 100nM 2-mercaptoethanol (Thermo Fisher Scientific), 3µM CHIR99201, 1µM PD032552 inhibitors (both from Cell Guidance Systems) and 1000 U/mL units of Leukemia Inhibitory Factor (Merck). Cells were passaged with 0.05% accutase (Thermo Fisher Scientific) every two days and maintained at 37°C with 8% CO2 in a humidified chamber. For *in vitro* differentiation, 10^6^ mESCs were grown in 10 cm bacteriological Petri dish with differentiation medium containing 25% Advanced DMEM, 25% DMEM/F12 GlutaMAX, 50% Neurobasal medium, 1% of penicilline/streptavidin, 1x β-mercatoethanol and 11% of knock out serum, at 37°C with 5% CO2 (all products from Thermo Fisher Scientific). The differentiation medium was changed every day and embryoid bodies were collected by centrifugation for 4 minutes at 700 rpm and resuspended in 10ml of fresh medium.

Lymphoblastoid cell lines (LCLs) from healthy donors and ICF2 patients were cultured in RPMI 1640 supplemented with 15% FCS and antibiotics (Thermo Fisher Scientific) as described in (15).

### Generation of the Auxin-Induced-Degradation system of ZBTB24 in mESCs

*Os*TIR1-9xMyc sequence linked via cleavable P2A sequence was introduced by CRISPR/Cas9-mediated homologous recombination at the C-terminus of the essential ubiquitously expressed RCC1 protein (regulator of chromosome condensation 1) into Zbtb24^wt/mt^ mESCs [guide RNAs (gRNAs] are listed in **Table S9**, all plasmids were a kind gift from by Dr. Alexei Arnaoutov, NICHD, NIH). Transfections were all performed using Lipofectamin2000 (Thermo Fisher Scientific). ESCs were selected using 7 μg/ml blasticidin and single cell clones were isolated by limiting dilution and the *os*Tir1 bi-allelic insertion was validated by PCR, Sanger sequencing, and western blotting. Sequences encoding a minimal functional auxin-inducible degron (mAID) and HA-tag were introduced by CRISPR/Cas9-mediated homologous recombination to the N-terminus of ZBTB24 protein (gRNAs are listed in **Table S9** were cloned in pSpCas9(BB)-2A-GFP (PX458) (Addgene #48138), the plasmids containing HA tag, a micro-AID tag (71–114 amino-acid), and puromycin resistance were kind gift from Dr Alexey Arnaoutov (72). Following hygromycin selection at 150 μg/ml, single-cell clones were isolated by limiting dilution. These clones were then validated for the mAID insertion on WT allele, followed by the characterization of the AID system.

### RNA extraction and RT-qPCR analysis

Total RNAs from cell lines were isolated using TRI Reagent® (Merck) according to manufacturer’s instructions. Contaminant genomic DNA was eliminated with TURBO DNA-free Kit (Thermo Fisher Scientific). Reverse transcription was carried out using 500ng DNA-free RNA, 50μM random hexamers (Thermo Fisher Scientific), 10U of RNase inhibitor (New England Biolabs) and 200U of ProtoScript II Reverse Transcriptase (New England Biolabs). Complementary DNA reactions were used as templates for real-time quantitative PCR reactions (qPCR). qPCR was performed using the SensiFAST SYBR NoROX Kit (Bioline) supplemented with 0.2μM of specific primer pairs (**Table S9**) and run on a light cycler rapid thermal system (LightCycler®480 2.0 Real time PCR system, Roche) with 20 sec of denaturation at 95°C, 20 sec of annealing at 60°C and 20 sec of extension at 72°C for all primers and analyzed by the comparative CT(ΔCT) method using U6 RNA as an invariant RNA. Each data shown in RT-qPCR analysis is the result of at least three independent experiments.

### Protein extraction and western blot analysis

Cells were washed in PBS and collected. After centrifugation, cell pellets were resuspended in RIPA buffer (300 mM NaCl, 1% NP-40, 0.5% Na-deoxycholate, 0.1% SDS, 50 mM Tris pH 8.0, complete protease inhibitor cocktail (Merck)). Samples were incubated for 30min on ice and were then sonicated with a Bioruptor (Diagenode) at 4°C. Cell debris were removed by centrifugation for 10 min at 16,000g and supernatants were collected. For Western blot analysis, proteins were resolved by SDS–PAGE, transferred onto PVDF membrane (Thermo Fischer Scientific), incubated with the appropriate primary antibody (**Table S10**) and then with horseradish peroxidase-conjugated secondary antibodies. Protein detection was performed with ECL reagents (Thermo Fisher Scientific).

### Immunofluorescence

Cells were grown on glass coverslips until reaching 60% of confluence. Cells were then fixed with 4% paraformaldehyde in PBS (Electron Microscopy Sciences). After permeabilization with 0.1% Triton X-100 in PBS, samples were blocked in 2% donkey serum (Jackson Immunoresearch) in PBS followed by incubation with primary antibodies (**Table S10**) in blocking buffer overnight at 4°C. After incubation with secondary antibodies conjugated to Alexa-Fluor 488 or 594 (Jackson ImmunoResearch), coverslips were mounted in Vectashield medium containing 2μg/ml DAPI (Vector Laboratories). Image acquisition was performed at room temperature on a fluorescence microscope (Axioplan 2; Zeiss) with a Plan-Neofluar 100X/1.3 NA oil immersion objective (Zeiss) using a digital cooled camera (CoolSNAP fx; Photometrics) and METAMORPH 7.04 software (Roper Scientific, Trenton, NJ). Images presented correspond to one focal plane.

### Chromatin Immunoprecipitation (ChIP)

Mouse ESCs and LCLs were crosslinked with 1% formaldehyde for 10min at room temperature (RT) and the reaction was quenched by adding glycine at a final concentration of 0.125M for 5min at RT. Fixed cells were washed and adherent cells were harvested with phosphate-buffered saline (PBS). Cell nuclei were isolated using a cell lysis buffer (5mM Pipes pH8.0, 85mM KCl, 0.5% NP40, Complete protease inhibitor cocktail (Merck)) for 20min, at 4°C. Nuclei were then pelleted by centrifugation and lysed in a nuclei lysis buffer (50mM Tris HCl pH8.1, 150mM NaCl, 10mM EDTA, 1% SDS, Complete protease inhibitor cocktail (Merck)) for 20min, 4°C. Chromatin was subjected to sonication with a Bioruptor (Diagenode) yielding genomic DNA fragments with a bulk size of 150-300 bp. Supernatant was diluted 10 times in IP dilution buffer (16.7 mM Tris pH8, 167 mM NaCl, 1.2 mM EDTA, 1.1% Triton X-100, 0.01% SDS, Complete protease inhibitor cocktail (Merck)). Immunoprecipitations were performed overnight at 4°C on 40μg of chromatin with antibodies pre-bound to Dynabeads protein G (Thermo Fisher Scientific). The antibodies used are listed in **Table S10**. The Beads were then washed once with Low salt buffer (0.1% SDS, 1% Triton, 2 mM EDTA, 20 mM Tris pH 8, 150 mM NaCl), once with High salt buffer (0.1% SDS, 1% Triton, 2 mM EDTA, 20 mM Tris pH 8; 500 mM NaCl), once with LiCl wash buffer (10 mM Tris pH 8.0, 1% Na-deoxycholate, 1% NP-40, 250 mM LiCl, 1 mM EDTA) and twice with TE (10mM Tris pH 8.0, 1mM EDTA pH 8.0). The elution of chromatin was performed in Elution buffer (50mM Tris pH 8.0, 10mM EDTA pH 8.0, 1% SDS) at 65°C for 45min and crosslink was reversed O/N at 65°C after addition of NaCl to 0.2M final concentration. The eluted material was digested with 40μg of Proteinase K (Thermo Fisher Scientific) at 65°C for 2h. DNA was purified by phenol– chlorofom extraction and ethanol precipitation, resuspended in TE. Quantitative PCRs were performed using the SensiFAST SYBR NoROX Kit (Bioline) and analyzed on a light cycler rapid thermal system (LightCycler®480 2.0 Real time PCR system, Roche). ChIP-qPCR results were normalized on input signal (% input). As a negative control, an additional mock ChIP using IgG was systematically added to each experiment. Sequences of primers are listed in **Table S9**.

### ChIP-seq experiment and analysis

ChIP-seq libraries were prepared either from ZBTB24 ChIP, IgG and input samples in LCLs derived from healthy subjects (n=3) or from ZBTB24 ChIP and input samples in *Zbtb24^wt/wt^* mESCs (n=2) cultured in 2i conditions and at d3 of differentiation, using the MicroPlex Library Preparation Kit v2 (Diagenode) according to the manufacturer’s protocol. In brief, 1 to 25 ng were used as input material. After adapters ligation, fragments were amplified with Illumina primers for 10 cycles. Libraries were purified using AMPure XP beads protocol (Beckman Coulter, Indianapolis, IN) and quantified using the Qubit fluorometer (Thermo Fisher Scientific). Libraries’ size distribution was assessed using the Bioanalyzer High Sensitivity DNA chip (Agilent Technologies). Libraries were normalized and pooled at 2nM and spike-in PhiX Control v3 (Illumina) was added. Sequencing was done using a NextSeq 500 instrument (Illumina) in paired-end mode (2×75 cycles). After sequencing, a primary analysis based on AOZAN software (ENS, Paris, France) (73) was applied to demultiplex and control the quality of the raw data (based on bcl2fastq v2.20 and FastQC modules / v0.11.5). Quality of raw data has been evaluated with FastQC (https://www.bioinformatics.babraham.ac.uk/projects/fastqc/). Poor quality sequences have been trimmed or removed with Trimmomatic (74) software to retain only good quality paired reads. Reads were mapped to the reference genomes (mm10/hg38) using Bowtie2 v2.3.4.3 (75) with the option -k 2. Then, peaks were called with Macs2 v2.2.4 (76) using the input as control, an FDR of 0.01 and keeping all the duplicates. Peaks from multiple replicates were combined by taking regions covered by at least 2 replicates. Peak annotation was performed with ChiPseeker v1.18.0 (77) and Bedtools intersect v2.28.0 was used to annotate peaks with repeated elements obtained on the repeatmasker website (http://www.repeatmasker.org), enhancers obtained on FANTOM5 (78) and genome segmentation from ENCODE data (79, 80). Finally, motif detection was performed with RSAT (81). The Gene ontology enrichment analysis was performed using the clusterProfiler R package. The significant GO categories were identified with Benjamini-Hochberg adjusted p-value (p.adjust) ≤ 0.05.

### Bioinformatic analysis of ZBTB24 binding sites in DNA repeats and HORs

ZBTB24 occupancy on DNA repeats was determined using RepEnrich2 (38) with the following protocol available in https://github.com/nerettilab/RepEnrich2. Briefly, the reads were first uniquely aligned with Bowtie2 (-k 1) on the mm10 or hg38 genomes. Single and multiple reads were then separated, and multiple reads were assigned to a FASTQ file. In parallel, annotation of the repeats was performed using RepeatMasker annotations of the repeated elements. Overlap of single reads with repeated elements was tested. Multiple reads were aligned against the repeat assembly using Bowtie2. Finally, the number of reads aligning against families of repeated elements was determined by combining the counts of uniquely mapping reads and multi-mapping reads (fractional counts). Read counts were normalized by the library size and the input. Read mapping to HOR arrays was carried out described in (40). The human ChIP_seq and RNA-seq FASTQ files were filtered against a library containing 6,455,351 sequences of 18 mers alpha satellite seed sequences (82). Reads that contained at least one 18mers were retrieved and mapped with Bowtie2 on a reference composed of 64 centromeric HOR array consensus sequences (83).

### RNA-seq experiment and analysis

RNA-seq was performed on independent clones of mESCs (2 clones of *Zbtb24^wt/wt^* cells and of *Zbtb24^mt/mt^*cells) and on LCLs derived from healthy (n=2) and ICF2 (n=2) subjects. RNA quality was verified using the Bioanalyzer RNA 6000 Nano kit (Agilent). The libraries were prepared following the TruSeq Stranded Total RNA protocol from Illumina, starting from 1 µg of high-quality total RNA (RIN > 7). Paired end (2 × 75) sequencing was performed on an Illumina NextSeq 500 platform, demultiplexing and quality control were performed with the AOZAN software (ENS, Paris, France) (73) (based on bcl2fastq v2.20 and FastQC modules / v0.11.5). The quality of raw data has been evaluated with FastQC (https://www.bioinformatics.babraham.ac.uk/projects/fastqc/). Poor quality sequences have been trimmed or removed with Trimmomatic (74) software to retain only good quality paired reads. Star v2.5.3a (84) has been used to align reads on reference genome (mm10/hg38) using standard options. Quantification of gene and isoform abundances has been done with rsem 1.2.28, prior to normalisation on library size with DESEq2 (85) bioconductor R package. Finally, differential analysis has been conducted with edgeR (86) bioconductor R package. Multiple hypothesis adjusted p-values were calculated with the Benjamini-Hochberg procedure to control FDR.

### DNA methylation of satellite repeats by Southern blot

Genomic DNA was extracted from mESCs and LCLs using the Monarch Genomic DNA Purification Kit (New England Biolabs) according to the manufacturer’s instructions. Genomic DNA from mESCs and LCLs (500 ng) was digested respectively with 20 units of HpaII or HhaI enzymes (New England Biolabs) for 16 h to analyze the DNA methylation patterns of centromeric minor and alpha satellite repeats, respectively. Genomic DNA from *Dnmt1/Dnmt3a/Dnmt3b* triple-knockout (TKO) mESCs (70) and from *Dnmt1* knock-out (*Dnmt1-/-*) mouse embryonic fibroblasts (87) were used as controls of low CpG methylated cellular contexts. The digested DNA fragments were separated by electrophoresis using 1% agarose gels and transferred overnight to Hybond-N+ membranes (GE Healthcare) in 20 × SSC. After ultraviolet crosslink, the membranes were pre-hybridized in 6 × SSC, 5 × Denhardt and 0.1% SDS and then hybridized with ^32^P-labelled minor satellite (5′-ACATTCGTTGGAAACGGGATTTGTAGAACAGTGTATATCAATGAGTTACAATGAG AAACAT-3′) or alpha satellite (5’-ATGTGTGCATTCAACTCACAGAGTTGAAC-3’). The membranes were washed 3 times in 6X SSC and 0.1%SDS at 37°C and then subjected to phosphorimaging using FLA 7000 phosphorimager (Fuji). The quantification of radiolabeled signals intensity was determined using ImageJ software and normalized by the sum of the total signal per lane.

### Methylome analysis by RRBS

RRBS libraries were prepared by MspI digestion from 50 to 100 ng genomic DNA44 and sequenced in paired-end 2 × 75 bp on an Illumina HiSeq4000 at Integragen SA (Evry, France). We trimmed RRBS reads to remove low-quality bases with Trim Galore v0.4.2 and aligned reads to the mm10 genome with BSMAP v2.74 (parameters -v 2 -w 100 -r 1 -x 400 -m 30 -D C-CGG -n 1). We calculated methylation scores using methratio.py in BSMAP v2.74 (parameters -z -u -g) and filtered CpGs covered by a minimum of eight reads. DMRs were identified using eDMR from the methylKit R package with a minimum of seven differentially methylated CpGs, a difference in methylation >20% and a q-value <0.001.

### Visualization of sequencing data

For ChIP-seq data visualization, the plots of the distribution of reads density across all transcription start sites RefSeq genes were generated using deepTools (https://deeptools.readthedocs.io/). Bar plots, dot plots, Hierarchical clustering representations, Heatmaps, violin plots, were generated using functions in R software environment for statistical computing. Samples clustering and heatmap visualizations were performed using the ComplexHeatmap and the gplots **R** packages (https://www.bioconductor.org/). For hierarchical clustering, the hclust function were used with “euclidean” and “ward.D2” method parameters respectively for distance and agglomeration. Basic stacked bar plots and Violin were build using the ggplot2 R package.

## DATA AVAILABILITY

The ChIP-seq, RNA-seq and RRBS data presented in this study have been deposited in the NCBI Gene Expression Omnibus (GEO) under the accession number GSEXXXXXX.

The following published datasets were used: GSE111689 (23) and GSE95040 (15).

## ACKNOWLEDGEMENTS

The authors would like to thank Drs Florence Larminat, Daniele Fachinetti, Catalina Salinas for insightful discussions about this work, Dr Riccardo Gamba for support and advises in centromeric data analysis, Laure Ferry and the Epigenomic Core Facility of the UMR7216 for help with pyrosequencing, Franck Letourneur and Juliette Harmoune of the sequencing platform core facility Genom’IC at Cochin Institute in Paris, Damien Ulveling and Justine Guegan of the bioinformatics and biostatistics core facility ICONICS in Paris for their help and support with sequencing data analysis. We acknowledge all the patients and their family members for their participation in this study.

## AUTHOR CONTRIBUTIONS

G.G., E.B., S.M., I.I., A.J., T.D. and G.V., performed the experiments. G.G., M.B, T.D., M.D., M.W. and G.V. performed bioinformatics analysis of sequencing data. G.V. and C.F. designed the study. G.V. and C.F. supervised the project. G.G., E.B. and G.V. wrote the original draft of the manuscript. G.V. and C.F. edited the manuscript.

---

**Figure S1.**
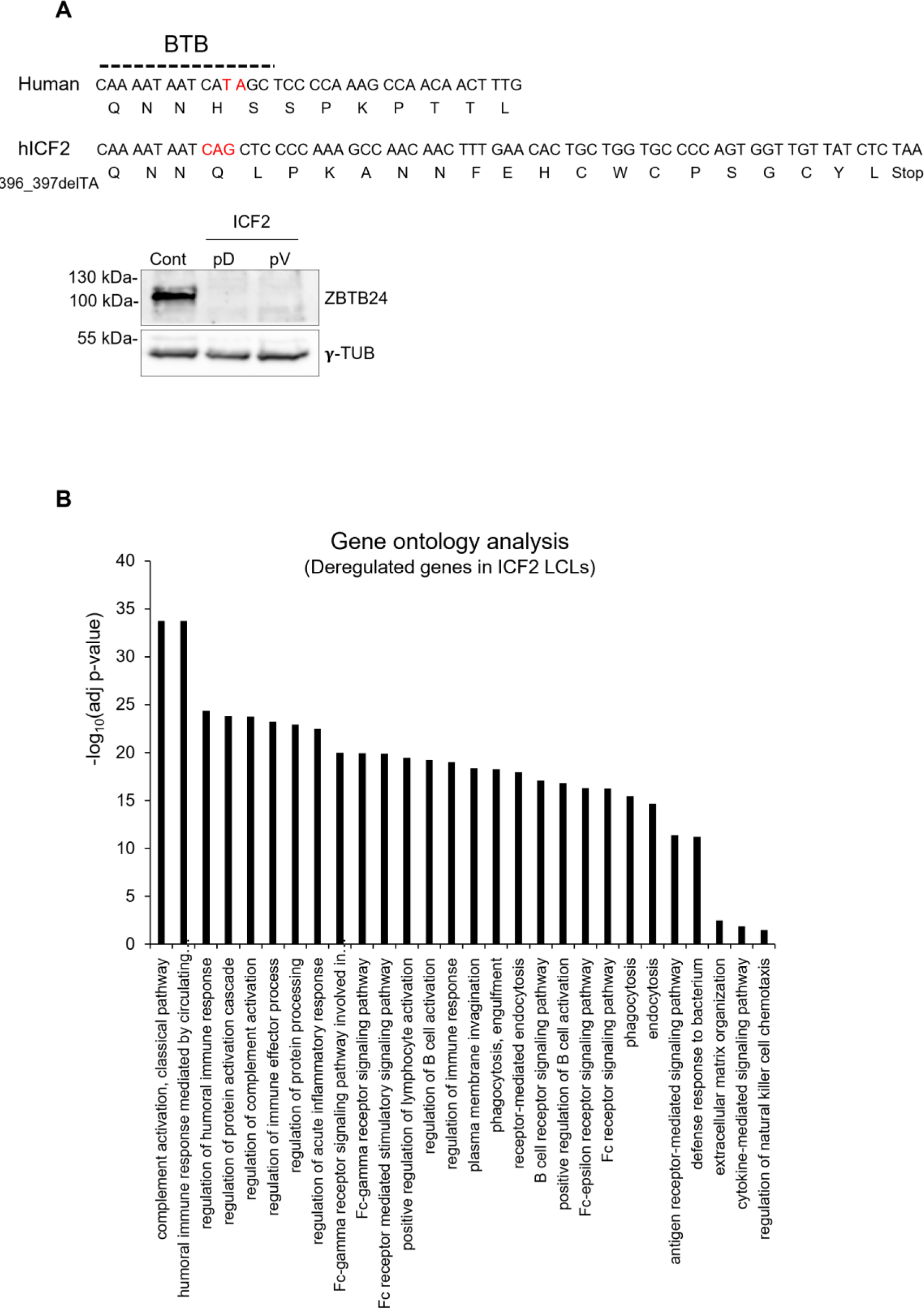
Analysis of gene expression alterations in ICF2 LCLs. (A) (top) Wild-type and mutant version of the human *ZBTB24* gene. The nucleotides mutated in ICF2 patients are indicated in red. The location of the BTB domain is indicated. (bottom) Western blot analysis of ZBTB24 protein levels in ICF2 patients (pD and PV) carrying the mutation shown above; γ-Tubulin is a loading control. (B) Gene ontology terms corresponding to deregulated genes in ICF2 patients’ LCL compared to healthy donors.

**Figure S2.**
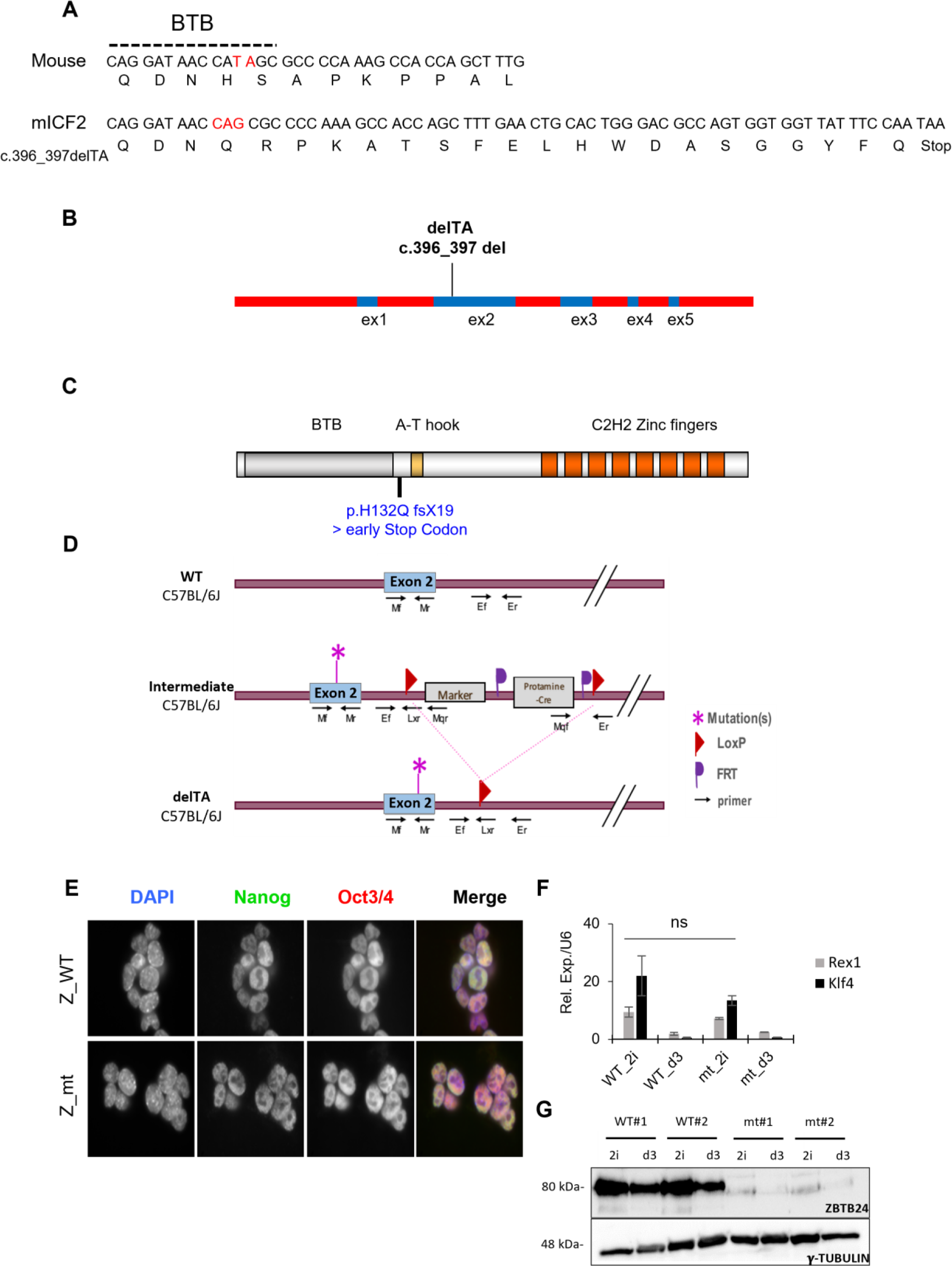
Analysis of the role of ZBTB24 during embryonic development. (A) Wild-type and mutant version of the mouse *Zbtb24* gene. The nucleotides mutated in the ICF2 mouse model presented in this study are indicated in red. The location of the BTB domain is indicated. (B) Schematic map of the *Zbtb24* mutant allele showing the position and the type of mutation. (C) Representation of ZBTB24 functional protein domains: BTB (Broad-Complex, Tramtrack and Bric a brac), AT-hook, and the eight C2H2 zinc finger domains. The position of the introduced mutation is indicated on the scheme. (D) Schematic representation of the WT and mutated *Zbtb24* alleles. The genetic background, the position of the loxP sites, the mutation, and the primers used for the genotyping of embryos are indicated. (E) Immunofluorescence showing the single cell expression of the pluripotency factors NANOG and OCT3/4 in *Zbtb24^wt/wt^* (Z_WT) and *Zbtb24^mt/mt^*(Z_mt) mESCs cultured in 2i medium. (F) Bar plots showing the relative expression levels of the pluripotency markers REX1 and KLF4, assessed by RT-qPCR and normalized to U6 small nuclear RNA. The graphs represent the mean of three experiments using WT and mutant (Mut) mESCs for ZBTB24 cultured in 2i medium and at d3 of differentiation. Error bars represent the standard error. n.s. non significant according to two-tailed t-test. (G) Western blot analysis of ZBTB24 protein levels in 2i condition and after three days of differentiation (d3) in *Zbtb24^wt/wt^*and *Zbtb24^mt/mt^* mESCs; γ-Tubulin is a loading control.

**Figure S3.**
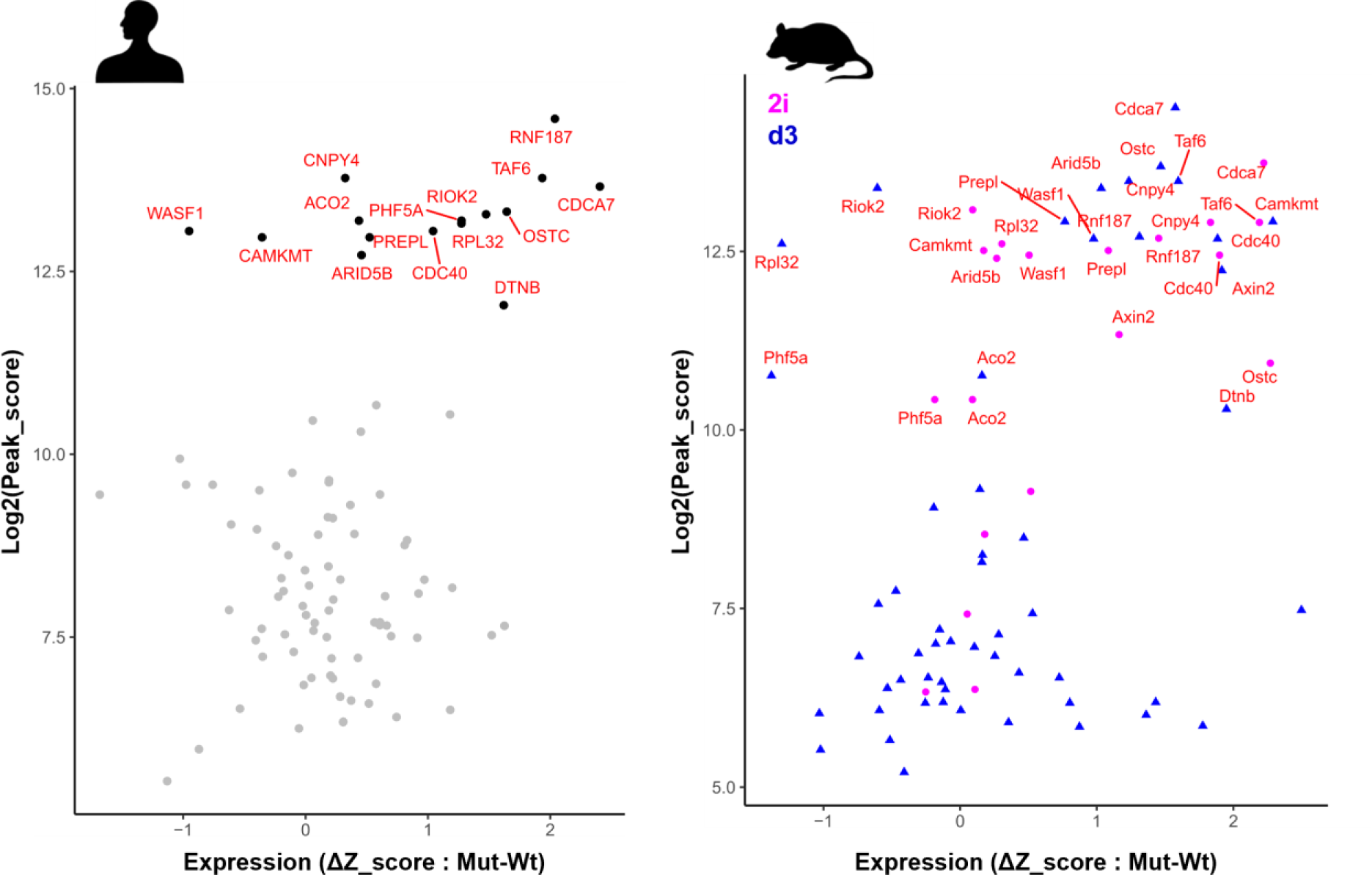
ZBTB24 targets the TSS and control the expression of a conserved set of genes between mouse and human. Scatter plots showing the distribution of the TSS of genes occupied by ZBTB24 according to their peak score (MACS2 score) and their expression status (ΔZ_score) in (left panel) LCLs mutant for ZBTB24 (Mut) compared to WT and in (right panel) *Zbtb24^-/-^* mES cells compared to *Zbtb24^+/+^* cultured in 2i medium (pink dots) or at day 3 of differentiation (d3, blue triangle points). The TSS of genes that are highly targeted by ZBTB24 and common to mouse and human are indicated by their gene symbol in red.

**Figure S4.**
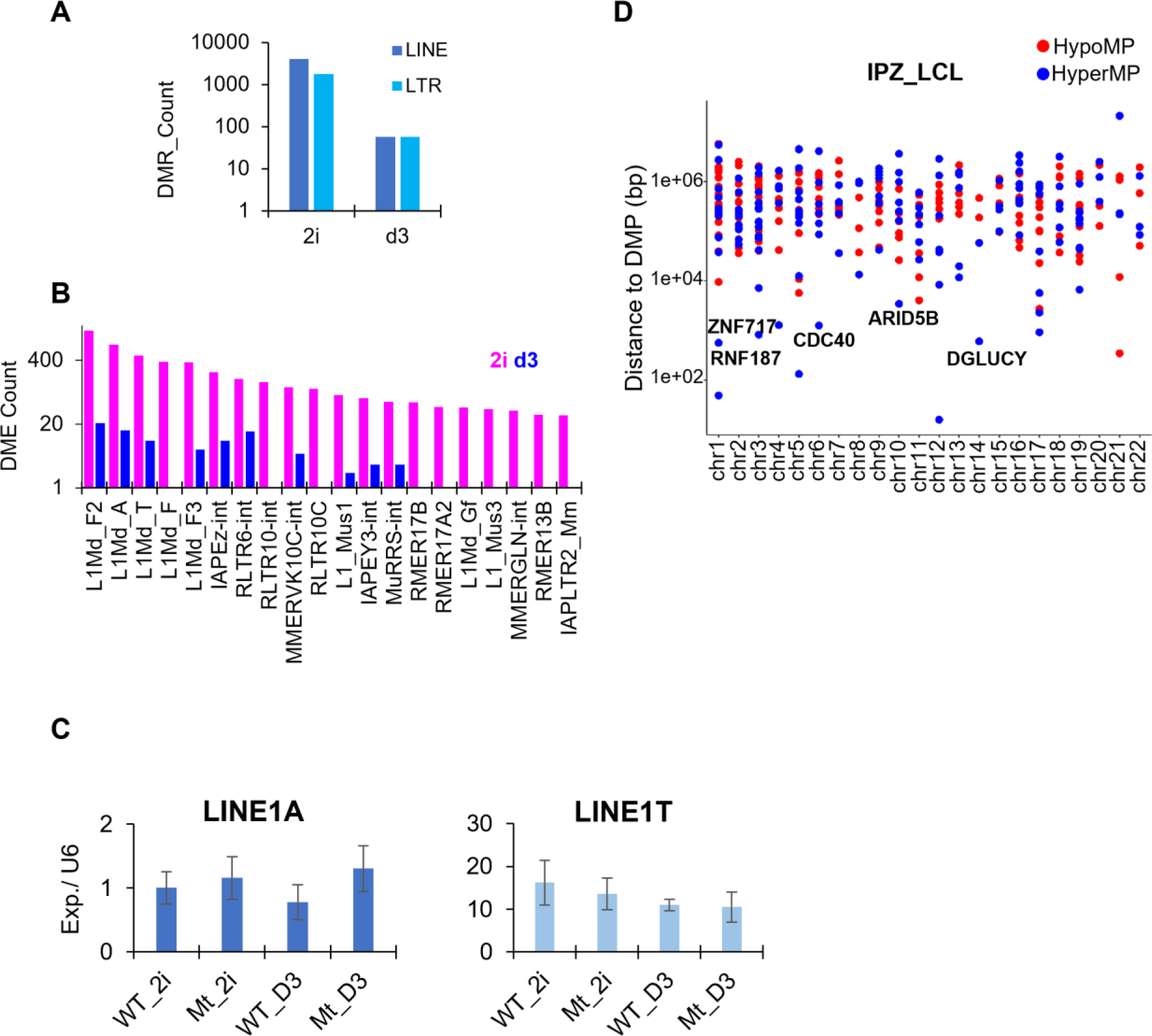
ZBTB24 LOF impacts DNAme of retrotransposon elements without impacting their transcriptional status. (A) Bar charts representing the number of the identified DMRs within LINE and LTR repeat class elements in mESCs. (B) Bar charts representing the number of differentially methylated elements (DME) within LINE and LTR repeat families in mESCs. (C) Bar plots showing the relative expression levels of the LINE1A and LINE1T elements, assessed by RT-qPCR and normalized to U6 small nuclear RNA. The graphs represent the mean of three experiments using WT and mutant (Mut) mESCs for ZBTB24 cultured in 2i medium and at d3 of differentiation. Error bars represent the standard error. (D) Dot plot showing the distance in base pairs (bp) between ZBTB24 peak summit identified in LCLs and Differentially Methylated Probes (DMP: |Δβ| ≥0.2, adjPval ≤0.05, hypomethylated sites in red, hypermethylated sites in blue) identified by HM450K bead chip technology in human ICF2 patient blood cells compared to healthy subjects. Dots which correspond to genes whose change in their level of expression is correlated to DMP are indicated by their symbol.

**Figure S5.**
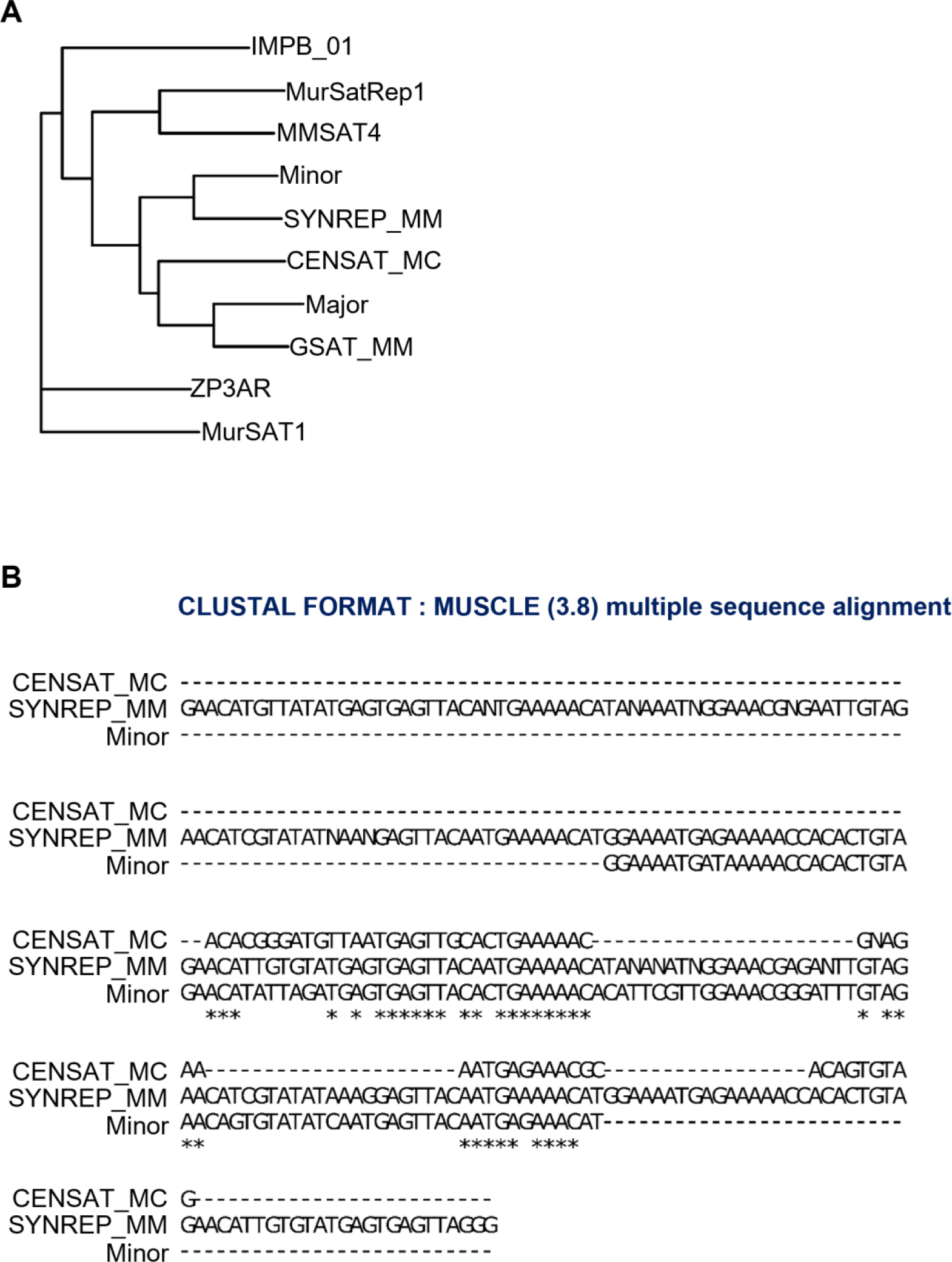
Repeatmasker annotation for minor satellite repeats. (A) Phylogenetic tree of satellite sequences in the mouse genome including the minor and major satellite sequences. Satellite sequences were retrieved from Dfam (https://dfam.org). Minor (120bp) and Major (234bp) satellite repeat unit sequences were respectively taken from Wong and Rattner (88) and From Manuelidis (89). (B) Comparison of Minor satellite sequence with CENSAT_MC and SYNREP_MM using MUltiple Sequence Comparison by Log-Expectation (MUSCLE).

**Figure S6.**
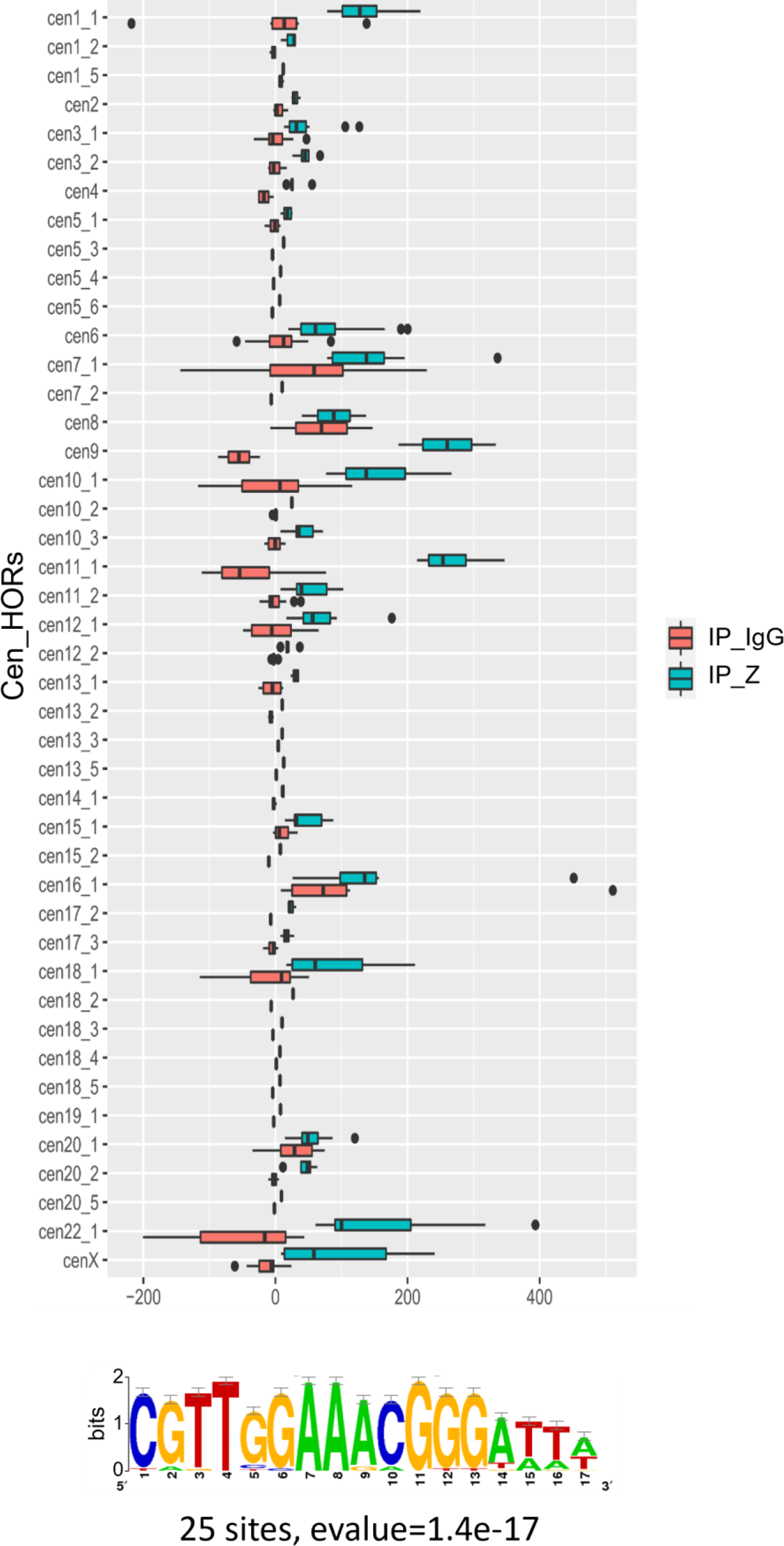
ZBTB24 is enriched at centromeric HORs. (Top) Box plot showing the enrichment of ZBTB24 relative to IgG within each centromeric HOR. The x axis shows the read density in ChIP-seq of ZBTB24 (IP-Z) and of IgG (IP_IgG) within each HOR. (Bottom) Predicted motif using Regulatory Sequence Analysis Tools (RSAT, http://rsat.sb-roscoff.fr/) based on ZBTB24 peaks within Cen_HORs is indicated below the box plot.

**Figure S7.**
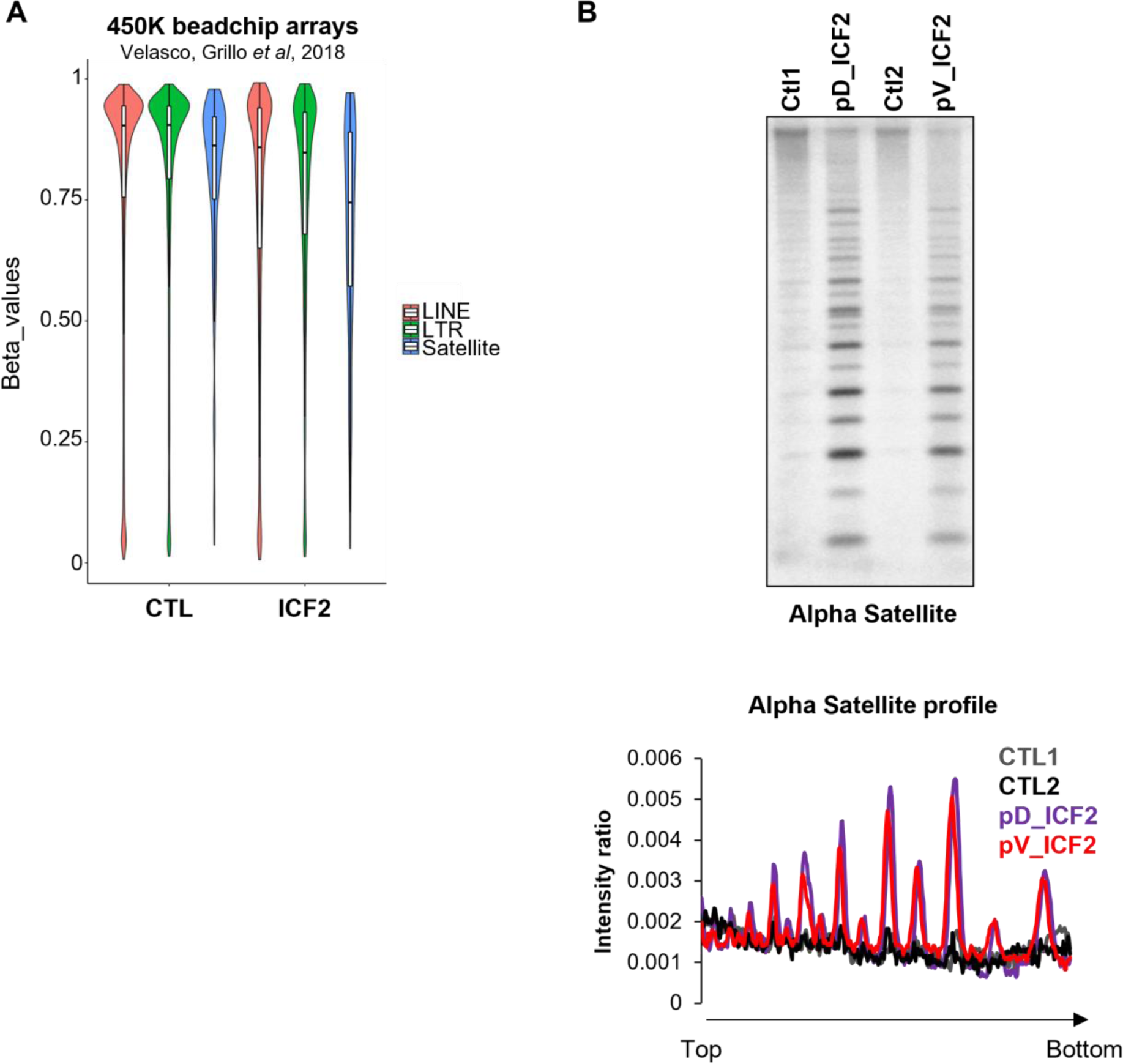
Centromeric satellite repeats are hypomethylated in human ICF cells. (A) Violin plot showing the normalized beta values (BMIQ normalization) of CpG probes of the Illumina Infinium methylation 450K (HM450K) bead chip technology, annotated in LINE, LTR and Satellite repeats on autosomes (n=22,821 probes). The data were generated from DNA of human primary blood cells derived from healthy subjects and ICF2 patients (15). (B) Southern blot and density plot of Alpha satellites repeats obtained from the digestion of genomic DNA of LCLs from healthy (CTL1 and CTL2) and ICF2 subjects (pD and pV) by the methylation-sensitive enzyme HpyCH4IV. The plot represents the profile of the radiolabeled signals intensity detected on each lane of the Southern blot after hybridization of radiolabeled alpha satellite probes and normalized by the sum of the total signal per lane (from the top toward the bottom of the lane).

**Figure S8.**
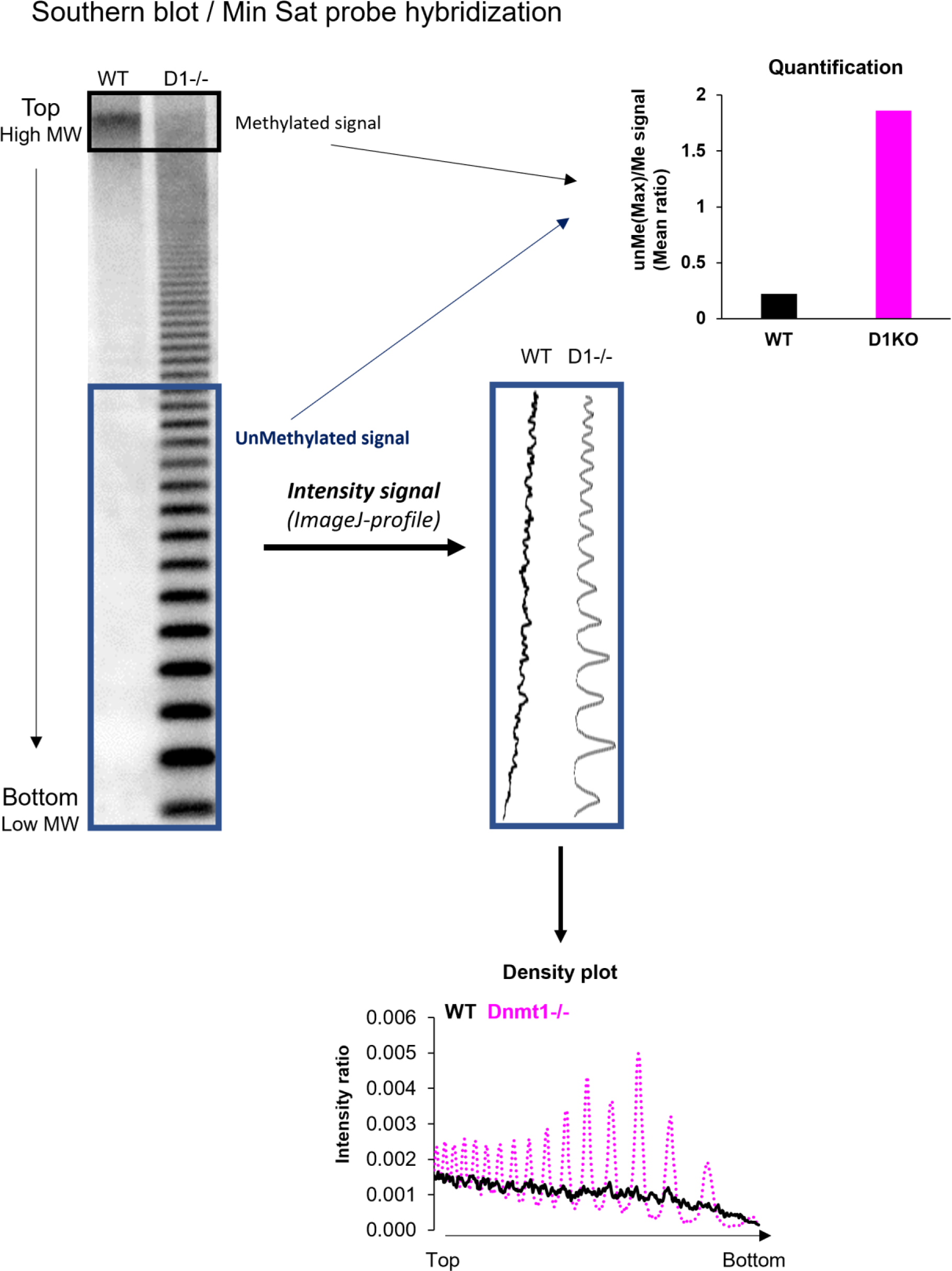
Representation of southern blot by density plots and quantification of DNA methylation signals. An example of how the density and the quantification plots were generated from the Southern blots is depicted in this scheme. To ease the comprehension, we used a Southern blot (top left) showing a HpaII digestion profile of minor satellite repeated DNA from a methylated (mouse embryonic fibroblast WT) and unmethylated context (Mouse Embryonic Fibroblast MEF *Dnmt1^-/-^*, DNMT1 being the DNA methyltransferase ensuring the maintenance of CpG methylation profiles through cell division). *Dnmt1^-/-^* MEF cells were kindly provided by Howard Cedar (87). The presence of a ladder with sharp bands in the *Dnmt1^-/-^* derived sample indicates that HpaII enzyme was able to cut DNA in the interrogated genomic region (minor satellite sequences), thus showing DNA hypomethylation of these sequences. The density plots (bottom) were generated from the intensity ratio of the radiolabeled signals of the Southern blot determined by using ImageJ software and normalized by the sum of the total signal per lane. The quantification of the DNA methylation (top right, bar plot) signal was calculated by the ratio of the mean of the unmethylated signal (unMe, represented as a ladder on the Southern blot) and of the methylated signal (Me, represented as the signal with the highest molecular weight MW at the top of the blot).

**Figure S9.**
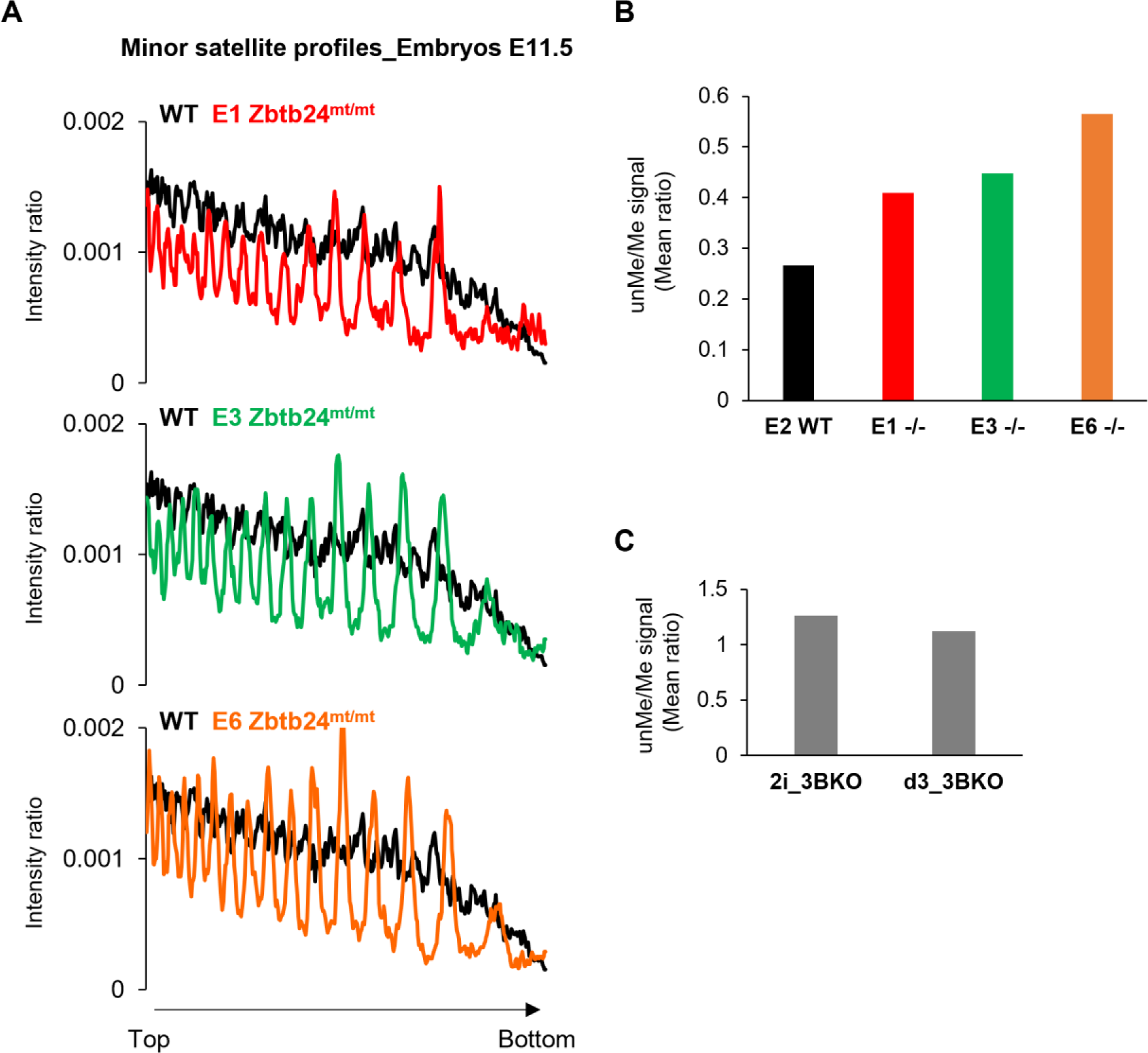
Centromeric satellite repeats are hypomethylated in mouse embryos mutant for ZBTB24. (A) Density plot of Southern blot band profiles obtained from the digestion of genomic DNA of WT and mutant embryos for ZBTB24 (*Zbtb24^mt/mt^*) at 11.5 dpc by the methylation-sensitive enzyme HpaII. The plot represents the profile of the radiolabeled signals intensity detected on each lane of the Southern blot after hybridization by radiolabeled minor satellite probes and normalized by the sum of the total signal per lane (from the top toward the bottom of the lane). (B) Quantification of DNA methylation changes at minor satellite in WT and mutant embryos for ZBTB24 (*Zbtb24^mt/mt^*) at 11.5 dpc. (C) Quantification of DNA methylation changes at minor satellite in *Dnmt3b^mt/mt^* mESCs (3BKO) cultured in 2i medium or after 3 days of differentiation.

**Figure S10.**
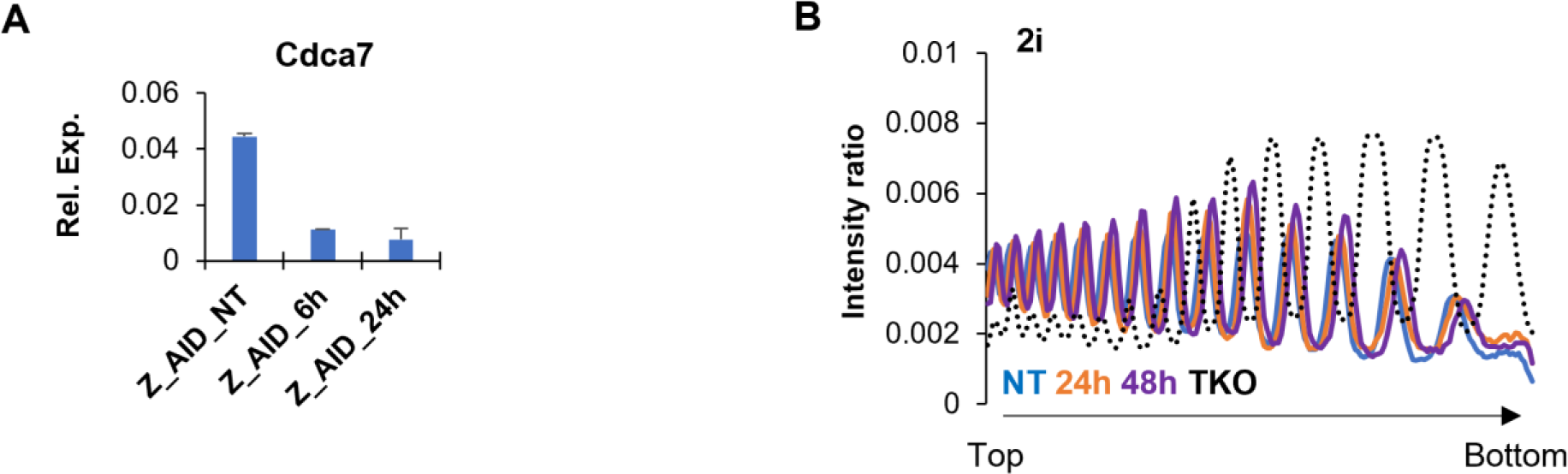
Validation of the AID-degron system for ZBTB24. (A) Expression level of *Cdca7* in mESCs AID_ZBTB24 lines cultured in 2i medium, treated or not (Z_AID_NT) for 6H (Z_AID_6h) or 24H (Z_AID_24h) with auxin. RT-qPCR experiments were performed and presented as a relative expression to U6 snRNA levels. Error bars represent standard error (n=3 independent experiments on two different clones). (B) Density plot of southern blot band profiles obtained from the digestion of genomic DNA of mESC AID_ZBTB24 lines treated or not (NT) for 24 h or 48 h with auxin and cultured in 2i medium. The plot represents the profile of the radiolabeled signals intensity detected on each lane of the Southern blot after hybridization by radiolabeled minor satellite probes and normalized by the sum of the total signal per lane (from the top toward the bottom of the lane). The DNA from mESCs triple knocked-out (TKO) for the DNMTs enzymes was used as a control of the hypomethylated context.

